# Mavacamten inhibits myosin activity by stabilising the myosin interacting-heads motif and stalling motor force generation

**DOI:** 10.1101/2025.02.12.637875

**Authors:** Sean N. McMillan, Jaime R. T. Pitts, Bipasha Barua, Donald A. Winkelmann, Charlotte A. Scarff

## Abstract

Most sudden cardiac deaths in young people arise from hypertrophic cardiomyopathy, a genetic disease of the heart muscle, with many causative mutations found in the molecular motor beta-cardiac myosin that drives contraction. Therapeutic intervention has until recently been limited to symptomatic relief or invasive procedures. However, small molecule modulators of cardiac myosin are promising therapeutic options to target disease progression. Mavacamten is the first example to gain FDA approval but its molecular mode of action remains unclear, limiting our understanding of its functional effects in disease. To better understand this, we solved the cryoEM structures of beta-cardiac heavy meromyosin in three ADP.Pi-bound states, the primed motor domain in the presence and absence of mavacamten, and the sequestered autoinhibited interacting-heads motif (IHM) in complex with mavacamten, to 2.9 Å, 3.4 Å and 3.7 Å global resolution respectively. Together with quantitative crosslinking mass spectrometric analysis, these structures reveal how mavacamten inhibits myosin. Mavacamten stabilises ADP.Pi binding, stalling the motor domain in a primed state, reducing motor dynamics required for actin-binding cleft closure, and slowing progression through the force generation cycle. Within the two-headed myosin molecule, these effects are propagated and lead to stabilisation of the IHM, through increased contacts at the motor-motor interface. Critically, while mavacamten treatment can thus rescue cardiac muscle relaxation in diastole, it can also reduce contractile output in systole in the heart.

## Introduction

Hypertrophic cardiomyopathy (HCM) affects at least 1 in 500 people and is the most common cause of heart failure in the young^1^. It is a genetic disease of the sarcomere, with ∼40 % of known disease-causing mutations found in the beta-cardiac myosin (βCM) heavy chain gene (MYH7) ^2^ and causing ∼30% of disease cases^3^. The exact disease mechanism(s) for HCM remains unclear. Treatment typically involves symptomatic relief, with beta or calcium channel blockers, or invasive procedures, such as septal myectomy, which do not address the root cause of disease^4^. More recently, small molecule treatments have been developed to directly modulate βCM force production^5^ and tackle disease progression. Mavacamten^6,7^ is the first of these to be FDA approved, with many others in clinical trials^8,9^. However, our understanding of the molecular mechanism of mavacamten is limited, restricting our understanding of its functional effects within disease.

βCM is the molecular motor responsible for force generation in cardiac ventricular tissue^10^, fuelled by ATP and through its interaction with filamentous actin. A single βCM molecule is comprised of two heavy chains and four light chains, of which two are essential (ELC) and two regulatory (RLC). Each heavy chain consists of an N-terminal globular motor domain, light chain binding domain (LCD) (where one ELC and one RLC bind) followed by an alpha helical region through which two heavy chains dimerise to form a coiled coil. The coiled coil is further divided into subfragment 2 (S2) and the filament-forming light meromyosin (LMM). Each motor domain is divided into four subdomains, the N-terminal domain, lower 50k domain (L50), upper 50k domain (U50) and the converter that together with the LCD forms the lever^11,12^.

Myosin molecules form bipolar filaments, with their LMM tails in the filament backbone and their paired motor domains on the filament surface^13,14^. To drive contraction they work in concert, interacting with actin to generate force transmitted by their levers in response to nucleotide and actin-binding state^12^. The motor domain together with the lever, collectively termed S1 or a myosin head, as well as cardiac heavy meromyosin (cHMM) (myosin lacking its LMM region), are competent to produce force and thus frequently studied to understand the myosin mechanochemical cycle.

Upon ATP binding a myosin head undergoes lever priming and ATP hydrolysis, to form a primed conformation^15,16^. In this ADP.Pi bound state myosin has an open actin-binding cleft between the U50 and L50 domains and can only interact with actin weakly. Conformational change within the motor leads to stronger actin-binding, cleft closure, Pi release and myosin lever swing (powerstroke), generating force^17,18^. ADP release from the motor results in a further, more minor, shift in lever position^19^. Re-binding of ATP opens the actin binding-cleft, disassociating the motor from actin and the cycle starts again^15^. Phosphate release is the rate limiting step for βCM in both the absence (basal turnover) and presence of actin^20,21^.

Within the heart, muscle contraction and thus force production are tightly regulated by two distinct mechanisms^22^. Actin thin filament regulation acts as an on-off switch, controlling when contraction can occur through intracellular calcium levels dictating when sites on actin are available for myosin binding^23^. Myosin thick filament regulation fine tunes the force output of individual thick filaments by controlling the number of motors available to produce force within the filament^24^. To do this, βCM can form a sequestered state outside of the force generation cycle called the interacting-heads motif (IHM).

The IHM forms through the asymmetric interaction of two ADP.Pi bound primed motors from the same molecule folding down against their S2 coiled coil^13,14^. Within the IHM one of the heads (termed the blocked head: BH) is blocked from interacting with actin as its actin-binding cleft is sequestered by the converter of the other head (termed free head: FH). In filaments, the IHM state is further stabilised through interactions with the thick filament backbone, titin and myosin-binding protein C^13,14^.

In HCM, the number of βCM molecules available to interact with actin and produce force are hypothesised to increase^25,26^, resulting in diminished relaxation and the diastolic dysfunction observed clinically^27^. Thus, HCM may potentially be treated by therapeutics which either inhibit myosin activity or increase IHM formation.

Mavacamten is a cardiac myosin allosteric inhibitor that inhibits sarcomeric force production, reducing cardiac contraction in animal models, isolated cells and muscle fibres^28^. It inhibits basal and actin-activated ATPase activity by inhibiting Pi release and stabilises an autoinhibited off-state^29^, reducing the number of functionally available motors to generate contractile force^28^. However, although mavacamten can stabilise an autoinhibited off-state^29^, the detailed structural nature of this state, the mechanism of this stabilisation, and how it inhibits Pi release are unclear^30^.

Several recent studies have begun to build on biochemical observations and unpick the structural mechanism of mavacamten. Three cryoEM structures have resolved the human cardiac thick filament, two in the presence^13,14^ and one in the absence of mavacamten^31^. These structures demonstrated that mavacamten stabilises the IHM but do not have sufficient resolution (resolutions ranging from ∼20 Å to 6 Å) to observe the underlying mechanism. Additionally, a recent crystal structure of a bovine S1 fragment complexed with ADP.BeFx and mavacamten showed the mavacamten binding site and that it restrains the lever^32^. The authors used molecular dynamic simulations to investigate how mavacamten inhibits myosin activity and proposed that mavacamten binding alters the L50 domain actin-binding interface to form a motor incompetent for force generation^32^. Yet this proposed mechanism does not explain how mavacamten inhibits Pi release.

Here, we performed a comprehensive analysis of the effects of mavacamten on cardiac myosin, from in vitro motility assays to structure determination by cryoEM and cross-linking studies using mass spectrometry. We show how mavacamten allosterically stabilises the IHM and inhibits Pi release from individual motor domains. We see no evidence for a significantly altered L50 domain structure or destabilisation of the L50 domain helix-loop-helix actin-binding interface upon mavacamten binding, as recently proposed^32^. In the context of the thick filament structures^13,14,31^, as well as recent observations made on the actin activation of myosin^18^, our data allow us to present a full structural mechanism for mavacamten inhibition of myosin.

## Results

### Mavacamten stabilises an off-pathway stalled ADP.Pi state

To ensure our cHMM construct was functionally active and inhibited by mavacamten, as expected, we tested its ability to drive actin filament movement by use of an in vitro filament gliding assay^33^ in comparison to a single-headed myosin construct (cS1) (Extended Data Fig. 2, Supplementary Table. 1). The mavacamten concentration required to reduce motility to 50% (IC_50_) was 0.14 µM for cHMM and ∼4-fold higher for cS1 (0.62 µM) (consistent with previous reports^28^). When filament movement was tracked over time in the presence of mavacamten, we observed that the number of moving filaments did not change for cS1 but decreased for cHMM in a dose dependant manner (Extended Data Fig. 2d, Supplementary Movie 1), consistent with the cHMM construct forming the IHM and reducing the number of motors available to interact with actin and produce movement.

To corroborate this, we examined the ratio of open head to IHM cHMM molecules in the presence and absence of mavacamten (Extended Data Fig. 3) using a negative stain EM assay. We found that mavacamten significantly (P=0.0002, determined by an unpaired two-tailed student’s T-test with respect to the βCM control) increased the percentage of cHMM molecules in the IHM when compared to cHMM alone.

To understand how mavacamten elicits these effects we used cryoEM to solve the structure of the cHMM motor domain in an open heads conformation, in the presence (MD_mava_) and absence of mavacamten (MD), and the cHMM IHM conformation in the presence of mavacamten (IHM_mava_) (see Methods, Extended Data Fig. 1 & Supplementary Fig. 1 & 2 respectively).

We pre-incubated cHMM with ATP to enable formation of open and IHM ADP.Pi states prior to the addition of mavacamten (or DMSO as control), followed by subsequent cross-linking with BS3 (see Methods). We then optimised cryoEM grid and vitrification conditions for imaging either cHMM molecules in the open state or in the IHM, which were then selected for downstream processing accordingly (see methods).

The MD_mava_ and MD were resolved to 2.9 Å and 3.4 Å global resolution respectively (Supplementary Fig. 1; Supplementary Table. 2). To interpret the cryoEM density maps we built pseudo-atomic models using a homology model and molecular dynamics driven flexible fitting^34^ (Supplementary Table. 3). Both maps showed density for MgADP.Pi and a primed lever with associated ELC density, confirming they were in a primed conformation (Extended Data Fig. 4, Fig. 1.)

**Figure 1.**
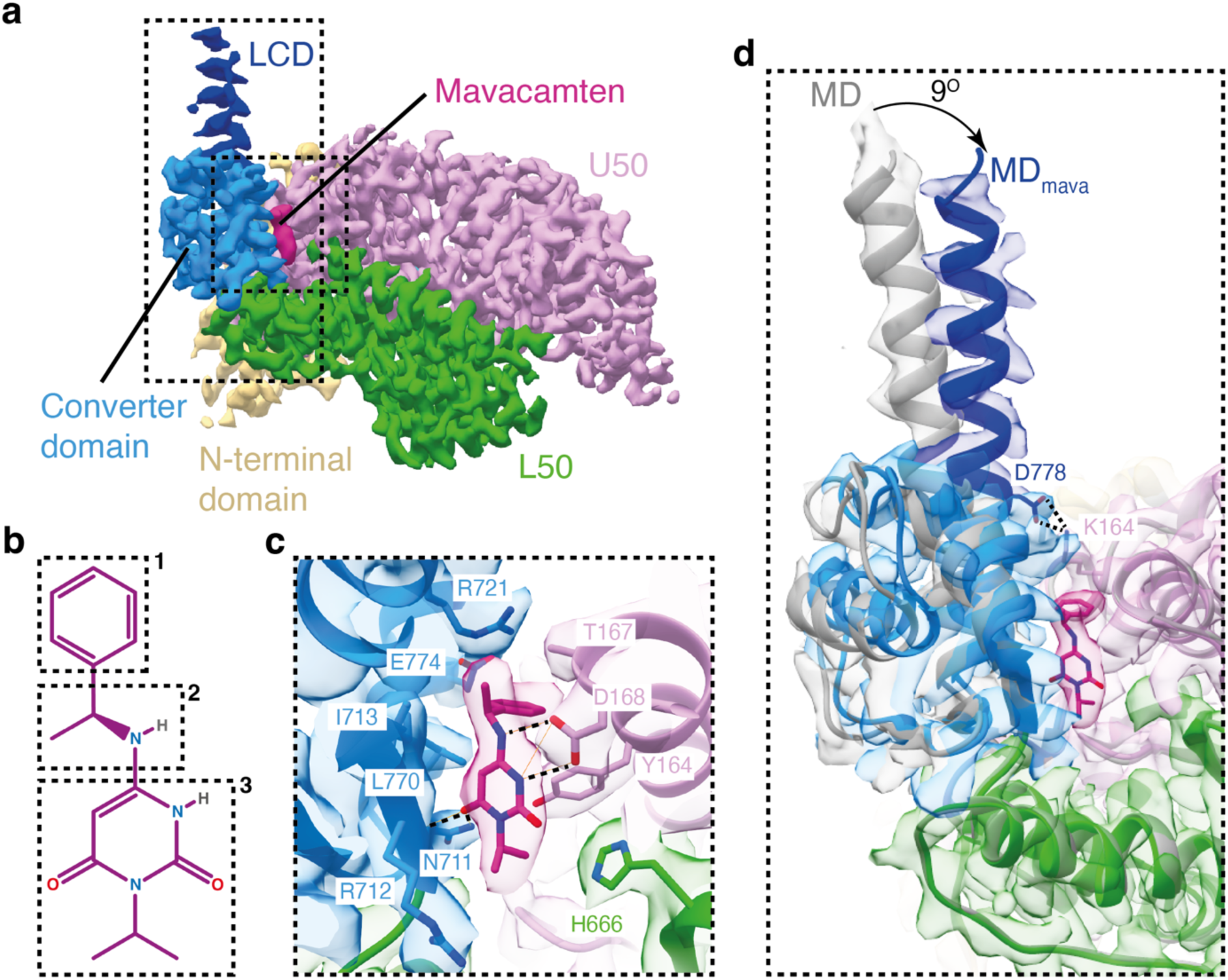
Mavacamten restrains the lever and D helix. (a) Segmented cryoEM map of the βCM motor in complex with mavacamten (MD_mava)_, split by subdomain (contour 0.6): N-terminal domain beige, L50 green, U50 pink, converter domain light blue, LCD dark blue and mavacamten in burgundy. (b) phenyl (1), methylethyl ester (2) and isopropyl pyrimidinedione (3) moieties of mavacamten (c) Mavacamten binding site highlighting key interactions: hydrophobic R721, L770, I713, H666 and T267; Ionic: Y164 and hydrogen bonding from N711, R712 and D168 (d) Overlay of MD grey and MD_mava_ coloured and segmented maps (contour MD: 0.5, MD_mava_: 0.6) showing 9° shift in the lever due to Mavacamten binding (global alignment).

Mavacamten was clearly resolved, positioned between the converter and the U50 (Fig. 1a). The mavacamten-protein interactions are predominantly hydrophobic formed by the sidechain backbones of residues R721, L770, and I713 on the converter with isopropyl pyrimidinedione, methylethyl ester and phenyl moieties of mavacamten (Fig. 1b-c) as well as H666 on the L50 and T167 on the U50 with the isopropyl pyrimidinedione and phenyl moieties. These hydrophobic contacts are then further supported by an ionic interaction between Y164 on the U50 and the isopropyl pyrimidinedione moiety and hydrogen bonding between N711, the backbone of R712 from the converter, and D168 from the U50 to the isopropyl pyrimidinedione and methylethyl ester moieties (Fig. 1b-c).

Binding of mavacamten rotates the lever 9° towards the U50 of the motor, perpendicular to the working stroke, when compared to the canonical primed MD conformation (Fig. 1d). This allows formation of a salt bridge between D778 and K146, creating additional communication between the lever and U50 domain (Fig. 1; Supplementary Movie. 2). Thus, mavacamten acts as a molecular glue, bonding the lever against the U50.

To understand how mavacamten may affect the association of myosin with actin in the weakly-bound ADP.Pi state, we aligned the MD and MD_mava_ structures on the L50 helix-loop-helix (HLH), the primary actomyosin binding interface^18^. Mavacamten binding subtly shifts the U50 relative to the L50, pinching the actin binding cleft in a motion distinct from cleft closure (Fig. 2a,b; Supplementary Movie. 2). This may be driven by the increased communication between the lever and the U50, which moves the D-helix towards the active site, altering the relative position of Y134 and nucleotide (Fig. 2c-e), compressing the ADP.Pi binding site.

**Figure 2.**
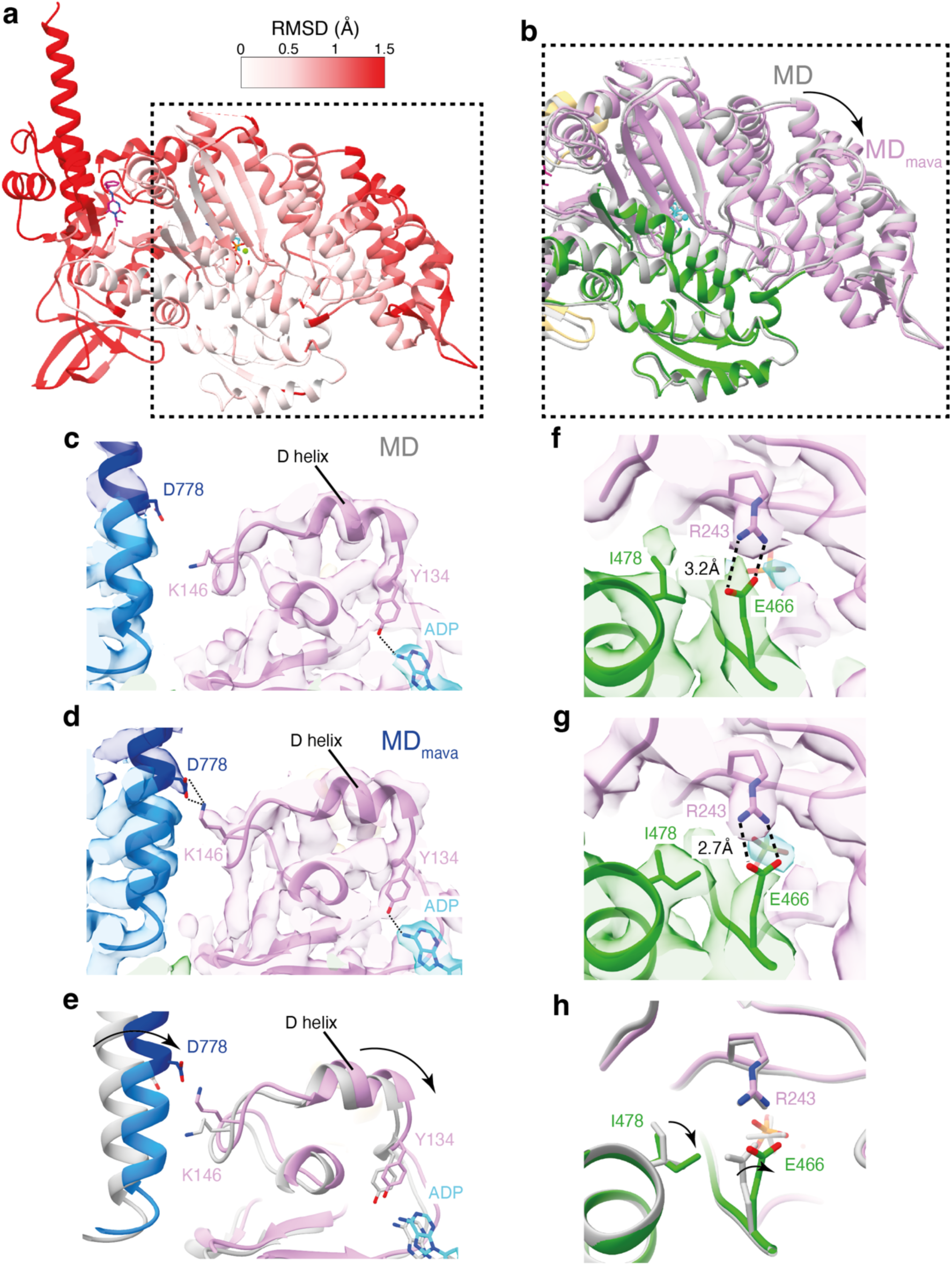
Mavacamten binding shifts the D-helix position stabilising Pi in the active site. (a) RMSD comparison between MD and MD_mava_ structures coloured on MD_mava_ model (aligned on the HLH). (b) Overlay of MD grey and MD_mava_ coloured, (aligned on the HLH) showing the subtle shift in the U50 (pink) towards the L50 (green). (c-e) Positioning of D778 and K146 in (c) MD, (d) MD_mava_ and (e) overlay MD gray and MD_mava_ coloured, highlighting communication between lever and D helix in MD_mava_. (f-h) MD and MD_mava_ structures in segmented maps respectively (contour MD: 0.6 and MD_mava_: 0.5), highlighting change in back door R243-E466 contact and I478. (h) Overlay of MD grey and MD_mava_ coloured highlighting change in residue positioning.

The shift of the U50 domain towards the L50 domain results in several rotamer changes around the backdoor, an ionic interaction between E466 and R243 that blocks the phosphate exit tunnel in the primed conformation^18^. I478 adopts a rotamer conformation facing the backdoor and the E466 side chain is rotated such that the distance between the R243 nitrogen (NH1) and E466 oxygen (OE1) is reduced from 3.2Å to 2.7Å (Fig. 2f-h; Supplementary Movie. 2). This shows that in the presence of mavacamten the back door interaction is stabilised, which may reduce the likelihood of Pi release.

These structural observations are not seen in the crystal structure of bovine S1 in complex with mavacamten (PDB ID: 8QYQ)^32^ Extended Data Fig. 5). Thus, our cryoEM structures demonstrate how mavacamten stabilises a stalled ADP.Pi state, and provide structural evidence for the mechanism of inhibited Pi release^29,32,35^.

It is important to note that the mavacamten induced structural changes are directionally opposed to those observed upon actin activation^18^, where the release of the Pi is accelerated ∼100-fold. Recent structural observations for myosin-5 have shown that upon initial binding of myosin to actin the U50 domain, particularly the D-helix, is cocked back towards the converter. This cocking back motion expands the nucleotide pocket destabilising the free Pi and promotes cleft closure^18^. Thus, we wondered whether mavacamten binding through cocking of the U50 forward, stabilising ADP.Pi binding, may also reduce motor domain dynamics to limit cleft closure. This would slow the transition from weak to strong binding of myosin to actin in the mechanochemical cycle, which is needed to sustain force and enable Pi release.

### Mavacamten binding reduces myosin lever and actin-binding cleft dynamics

To investigate the effect of mavacamten on primed motor lever and actin-binding cleft dynamics, we used a quantitative crosslinking mass spectrometry (qXL-MS) approach. We used the MS-cleavable, amine and hydroxyl reactive cross-linker, disuccinimidyl dibutyric urea (DSBU) to crosslink cS1 ADP.Pi in the absence and presence of mavacamten and then identified and quantified the crosslinks obtained using a label-free qXL-MS comparative approach^36–38^. 85 unique interpeptide cross-links and 49 monolinks changed in normalised signal intensity by at least two-fold between the two conditions (see Supplementary Material 1 & 2), indicating that adding mavacamten significantly affected cS1 conformational dynamics.

To interpret the effects of mavacamten on cS1 dynamics, we generated S1 models, S1_mava_ and S1 in the presence and absence of mavacamten, by superimposing the FH lever, ELC and RLC from our IHM_mava_ model (see Fig. 4a) onto our MD_mava_ and MD structures respectively. We then annotated the models with crosslinks that significantly increased or decreased in signal intensity 2-fold accordingly and measured their Cα-Cα distances (Supplementary Table 4, p<0.05 across three replicates). Crosslinks within the accepted DSBU reaction distance of 30 Å Cα-Cα were expected^39^. Comparatively, the observation of a crosslink between two residues with a Cα-Cα distance >30 Å indicated conformational dynamics that allowed the two reacting side chains to come into range.

Thus, if an increase in a crosslink signal intensity in the presence of mavacamten was observed with the same Cα-Cα distance in both S1_mava_ and S1 models, the rise in signal intensity indicated increased sampling of that conformational state and reduced dynamics. Whereas, an increase in signal intensity for a crosslink with a Cα-Cα distance >30 Å would suggest increased dynamics, as there was an increase in reactivity, e.g. increased time in which the two residues sampled a conformation outside our model in sufficient proximity to react.

To enable understanding of the changes in conformational dynamics that occur between individual motor subdomain domains, or the motor and LCs, on the addition of mavacamten, we focussed on interpreting interdomain crosslinks. 45 of the identified crosslinks were interdomain, of which we could annotate 29 on our models (Fig. 3a,b; Supplementary Table 4). The remainder could not be annotated as they were located within unmodelled or flexible regions, such as the N-terminal extension of the ELC or myosin loop2.

**Figure 3.**
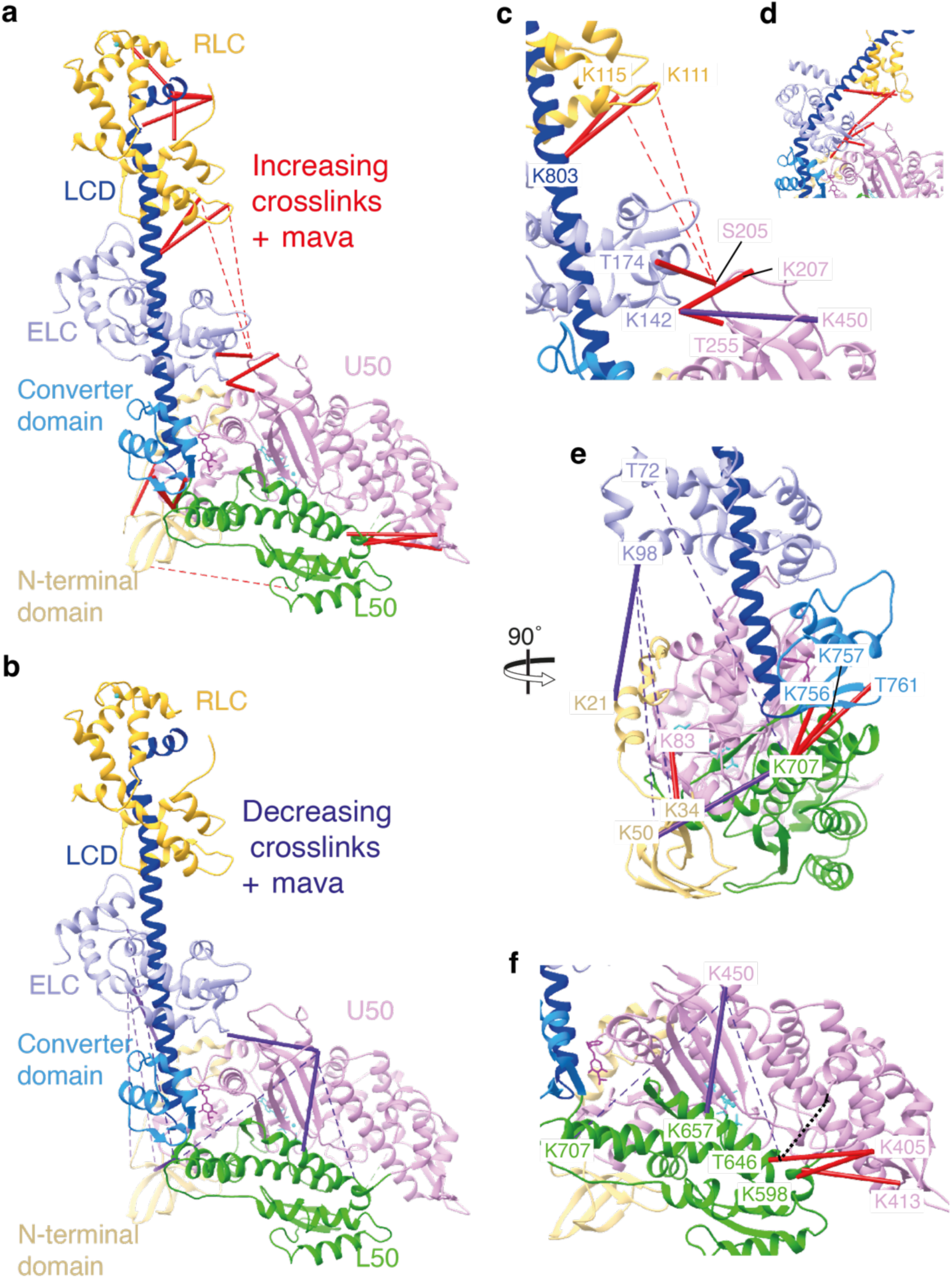
qXL-MS analysis of crosslinks in the presence of mavacamten. Overview of crosslinks observed to (a) increase and (b) decrease in the presence of mavacamten mapped onto the S1_mava_ model coloured by subdomain: N-terminal domain beige, L50 green, U50 pink, converter domain light blue, LCD dark blue, ELC light purple, RLC yellow and mavacamten burgundy. Solid lines denominate DSBU crosslinks with a <30 Å Cα-Cα distance whilst dashed lines represent crosslinks with a >30 Å Cα-Cα distance, red = increasing, purple = decreasing. (c-d) Magnified view of increasing crosslinks between the LCs and U50 subdomain, with a (c) open head and (d) BH lever position. (e) Overview of crosslinks showing increased stability of lever position in the presence of mavacamten (f) Overview of crosslinks showing changes in U50-L50 position in the presence of mavacamten. Loop2 is represented by a dotted black line. Crosslinks shown here are further described in Supplementary Table 4.

In the presence of mavacamten, the number of crosslinks (<30 Å Cα-Cα distance) increased significantly between residues that connect the U50 domain to the ELC (Fig. 3a,c; Supplementary Table 4). This is consistent with the lever being pulled in towards the U50 domain when mavacamten binds, as observed in our cryoEM structures (Fig. 1). Crosslinking was also increased between the RLC (K115 and K111), and the LCD (K803), and loop1 (S205). The RLC-loop1 crosslinks had a Cα-Cα distance >30 Å in our S1 models but would have a Cα-Cα distance <30 Å if the lever was bent at the pliant point, forming the IHM blocked head lever position (Fig. 3d). Thus, these crosslinks can also be interpreted to indicate increased proximity of lever position relative to the U50 domain in the presence of mavacamten (Supplementary Table 4). In the presence of mavacamten, the number of crosslinks between the base of the converter (CON) and the L50 domain (K707_L50_-K757_CON_, K707_L50_-T761_CON_) (Fig. 3e) increased significantly, without an increase in Cα-Cα distances in our S1 models, indicating increased stability of L50-converter position. Exploratory crosslinks (Cα-Cα distance >30 Å) between the ELC T72 and K98 and N-terminal domain (K21, K34 and K50) and converter (K707) respectively were significantly decreased in the presence of mavacamten (Fig. 3b,e). This suggests that the lever position is less dynamic, exploring a narrower range of primed conformations, preventing sampling of states that would allow these crosslinks to form.

When considering actin-binding cleft dynamics, the cleft is ordinarily thought to open and close rapidly on the µs timescale^16^ but with an equilibrium strongly favouring the open cleft position, and thus retention of Pi in the active site. In the presence of mavacamten, crosslinks (<30 Å Cα-Cα distance) between the HCM loop in the U50 domain (K405 and K413), and the strut (K598) and C-terminal end of loop2 (T646) within the L50 domain (Fig. 3f; Supplementary Table. 4), bridging the actin-binding cleft increase in signal intensity. This suggests that cleft dynamics are reduced when mavacamten binds, as there is an increase in sampling of conformations which enable these crosslinks to form. This reduction in cleft dynamics may help prevent the weak to strong binding transition, inhibiting phosphate release and effective actomyosin cross-bridging.

Crosslinking of loop2 (K633-K640) to loop2 K633 and L50 K657, also significantly increased, alongside a decrease in K657-K450 crosslinking in the presence of mavacamten. This suggested that the binding of mavacamten stabilises the conformation of loop2 enabling K657 to more readily crosslink with K633 than with K450. The number of exploratory crosslinks (>30 Å Cα-Cα distance) between U50 K450 and L50 K707 and T646 was reduced in the presence of mavacamten, indicating reduced motor domain dynamics due to interaction with the drug.

Strikingly, the single crosslink between K570-K572 within loop3 of the L50 had the largest increase in signal intensity in the presence of mavacamten (Supplementary Material 1). This is beautifully explained when we examine the hydrogen bonding network of loop 3 within the MD and MD_mava_ structures respectively (Extended Data Fig. 6). K570 is solvent exposed within both structures but in the presence of mavacamten, an ionic interaction between D469 and K572 is lost and instead D587 interacts with R567. This leaves K572 without a binding partner, making it available for crosslinking to K570 and accounting for this dramatic change in reactivity.

Although the rearrangement of this hydrogen bonding network may increase the dynamics of loop 3 in the presence of mavacamten, there is a corresponding decrease to the exploratory crosslink K565_L50_-K707_L50_, between the N-terminal end of loop3 and the base of the converter (Supplementary Material 1). This allows us to reasonably conclude that the localised increase in dynamics of loop 3 is due to allosteric communication and not a wholesale increase of dynamics within the L50. Corroborating this, we see no evidence within the cryoEM density maps for a decrease in stability of the HLH in the presence of mavacamten (Supplementary information Fig. 1l,m).

In addition to changes in crosslinking, we also observed changes to monolink signal intensity, where the crosslinker reacts with a residue and solvent. Monolinks for the ELC residues K142 and K147 and LCD residue K803 decrease in the presence of mavacamten as the ELC and LCD residues are instead able to crosslink with the U50 and RLC respectively (see Supplementary Material 1). The most significant decrease in monolinking in the presence of mavacamten was K146_U50_, which is expected given the formation of an ionic interaction between K146_U50_ and D778_LCD_ in the presence of mavacamten in our structures that increases communication between the lever and nucleotide binding site.

Together, our combined cryoEM and qXL-MS structural analysis suggests that mavacamten stabilises ADP.Pi binding and limits actin-binding cleft closure, required for the weak to strong binding transition, to inhibit phosphate release. Thus, mavacamten stabilises an off-pathway stalled myosin motor conformation that is unable to progress through the mechanochemical cycle.

### Mavacamten restrains the IHM through stabilisation of the free head motor domain

To understand how mavacamten stabilises the IHM we used cryoEM to solve the structure of cHMM IHM in complex with mavacamten (IHM_mava_) to a global resolution of 3.7Å (Supplementary information Fig. 2; Fig. 4a) and compared it to our MD_mava_ structure and to the mavacamten free folded-back state (IHM^40^) previously reported (PDB ID: 8ACT)^40^. The overall appearance of the IHM_mava_ is consistent with the IHM structure. However, mavacamten binding induces distinct conformational changes within the motor domain of the BH and FH respectively.

**Figure 4.**
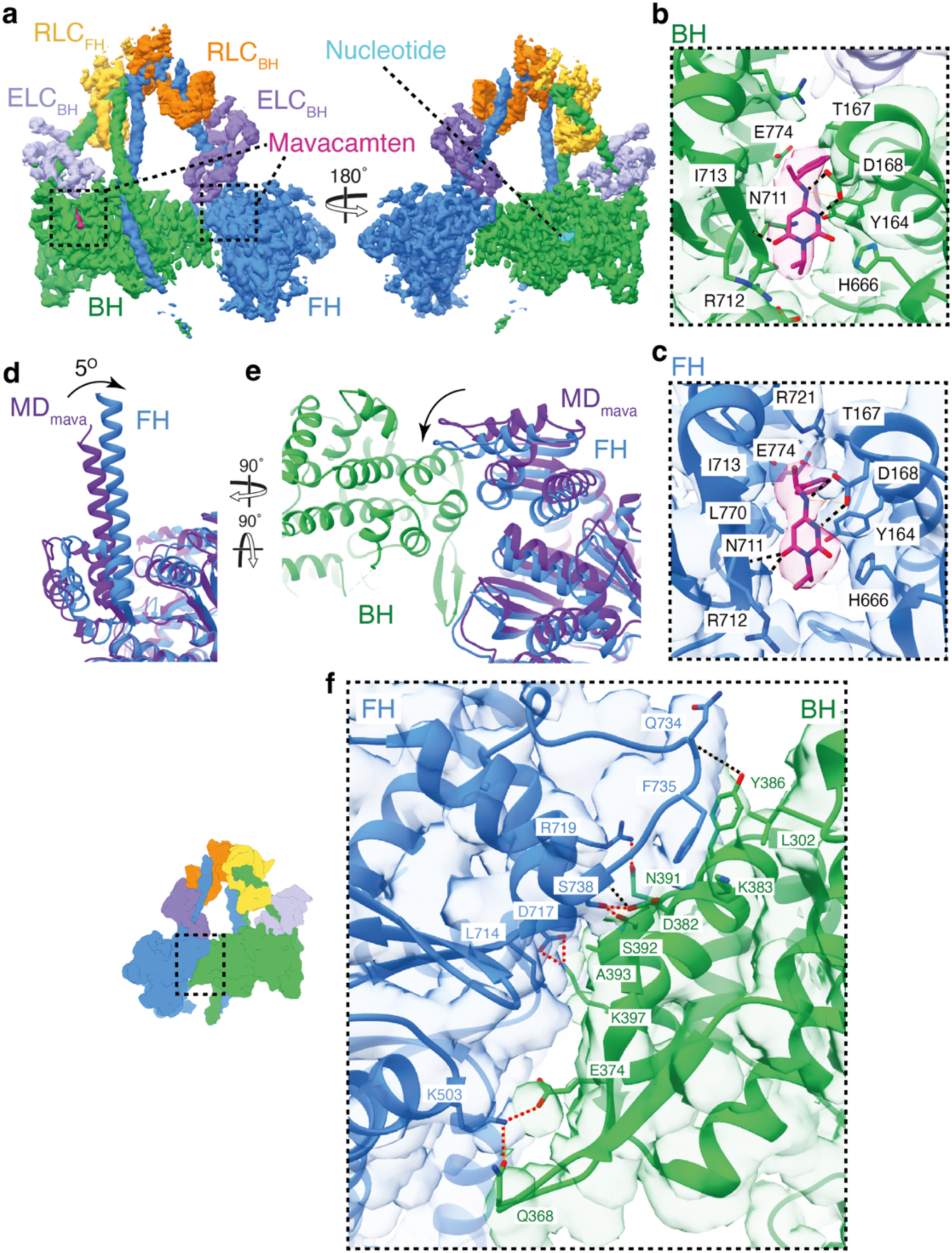
Mavacamten strengthens the motor-motor interface in the IHM by altering the FH lever conformation. (a) Segmented cryoEM map of the IHM_mava_ structure coloured by chain (contour: 0.08): BH green, FH blue, ELC_BH_ light purple, ELC_FH_ purple, RLC_BH_ yellow, RLC_FH_ orange, mavacamten burgundy, and the nucleotide in light blue. (b-c) The IHM_mava_ model fitted to the segmented map highlighting the BH and FH mavacamten pockets respectively. (d) Side view of the FH lever blue overlaid MD_mava_ dark purple (aligned on the motor D3-P710) highlighting the 5° shift of the lever. (e) Top-down view of (d) including the BH in green showing the movement of the free head converter. (f) Interaction interfaces between the U50_BH_-converter_FH_ (contour: 0.01), with H-bonds unique to the IHM_mava_ in red (compared to PDB ID: 8ACT^40^).

Mavacamten is bound to both the BH and FH of the IHM_mava_ (Fig. 4b,c). The FH mavacamten binding site is compressed, with the lever domain lying even closer to the U50 than in the MD_mava_ structure (Extended Data Fig. 7) yet maintains all interactions previously observed (Fig. 4c). Conversely, due to formation of the BH lever conformation, the mavacamten BH binding site is expanded (Extended Data Fig. 7) compared to the MD_mava_ structure, and its interaction with N711 and L770 is lost (Fig. 4b). This suggests mavacamten may have a greater effect on IHM_mava_ stabilisation through its interaction with the FH.

Indeed, the structural changes caused by mavacamten in the FH are the most prominent and provide a mechanism through which allosteric IHM stabilisation occurs. Comparison of the MD_mava_ and IHM_mava_ FH, by alignment on the main body of the motor (residues 3-710), shows the IHM_mava_ FH lever adopts a much sharper angle than in the MD_mava_ structure, rotating 5° perpendicular to the lever swing (Fig. 4d). If the same comparison is performed between the IHM^40^ FH and IHM_mava_ FH a rotation of 9° is observed in the same direction (Extended Data Fig. 8). The change in FH lever angle allows the converter to form a more extensive interface with the BH, strengthening existing contacts at the U50_BH_-converter_FH_ interface (Fig. 4e) between N391_BH_ and the backbone of S738_FH_ as well as Y386_BH_ and Q734_FH_ (Fig. 4f).

As the FH converter now packs more tightly against the U50_BH_ several new interactions form, supporting the interface. Interactions are formed between S738_FH_, S392_BH_ and D382_BH_, D717_FH_ and K397_BH_ as well as R735_FH_ and N391_BH_ in addition to a hydrophobic interaction between A393_BH_ and L714_FH_ (Fig. 4f; Supplementary Movie. 3). The movement also results in rearrangement of F735_FH_, which now forms a hydrophobic interface with K383_BH_, L302_BH_ and Y386_BH_. (Fig. 4f; Supplementary Movie. 3). Interactions are also altered at the HCM loop_BH_-transducer_FH_ interface with the formation of an ionic interaction between D409_BH_ and R249_FH_ (Supplementary Movie. 3). Finally, a new interaction interface between the U50_BH_ and the ELC_FH_ is able to form between K611_BH_ and D143_BH-ELC_ (Supplementary Movie. 3).

In summary, the change in FH lever angle upon mavacamten binding provides increased structural rigidity to the IHM through the generation of new interfaces, strengthening the motor-motor contact, providing a structural mechanism for its allosteric stabilising effects. This mechanism is supported by recent structural observations in which IHM crowns in a relaxed mavacamten-free thick filament structure were less ordered than in their mavacamten-bound counterparts, particularly in the free head of the disordered crown/crown 2^13,31^.

The blocked head conformation within the IHM_mava_ also deviates from that observed in both the MD_mava_ and the IHM^40^ structures but only in the relative positioning of the U50 domain. When the MD_mava_ is aligned on the L50_BH_ domain of the IHM_mava_ it is apparent that the U50_BH_ domain has moved across the L50_BH_ towards S2. If the same alignment is performed with the IHM^40^ and IHM_mava_ the same shift is observed (Extended Data Fig. 9). This conformational change in the U50_BH_ is needed to accommodate the altered converter_FH_ position upon mavacamten binding. Thus, the conformational changes observed in the blocked head upon mavacamten binding are a consequence of its IHM stabilising effect but not a contributor to IHM stabilisation.

Along with stabilising the motor-motor interface, mavacamten reduces the activity of the FH by restraining the nucleotide pocket. Comparison of the IHM_mava_ FH to our MD_mava_ structure and IHM^40^ shows the D helix_FH_, is moved further towards the nucleotide pocket in the IHM_mava_ structure (Fig. 5c-e; Extended Data Fig. 8). The U50_FH_ has also shifted relative to the L50_FH_ further pinching the actin binding cleft (Fig. 5a-b; Extended data Fig. 8). The resulting shift of the D-helix_FH_ and surrounding loops further reduces the available space for ADP dynamics, increasing the stability of ADP.Pi within the active site (Fig. 5f-g, Extended data Fig. 8e). Once again this is an opposite motion to that observed during actin activation^18^.

**Figure 5.**
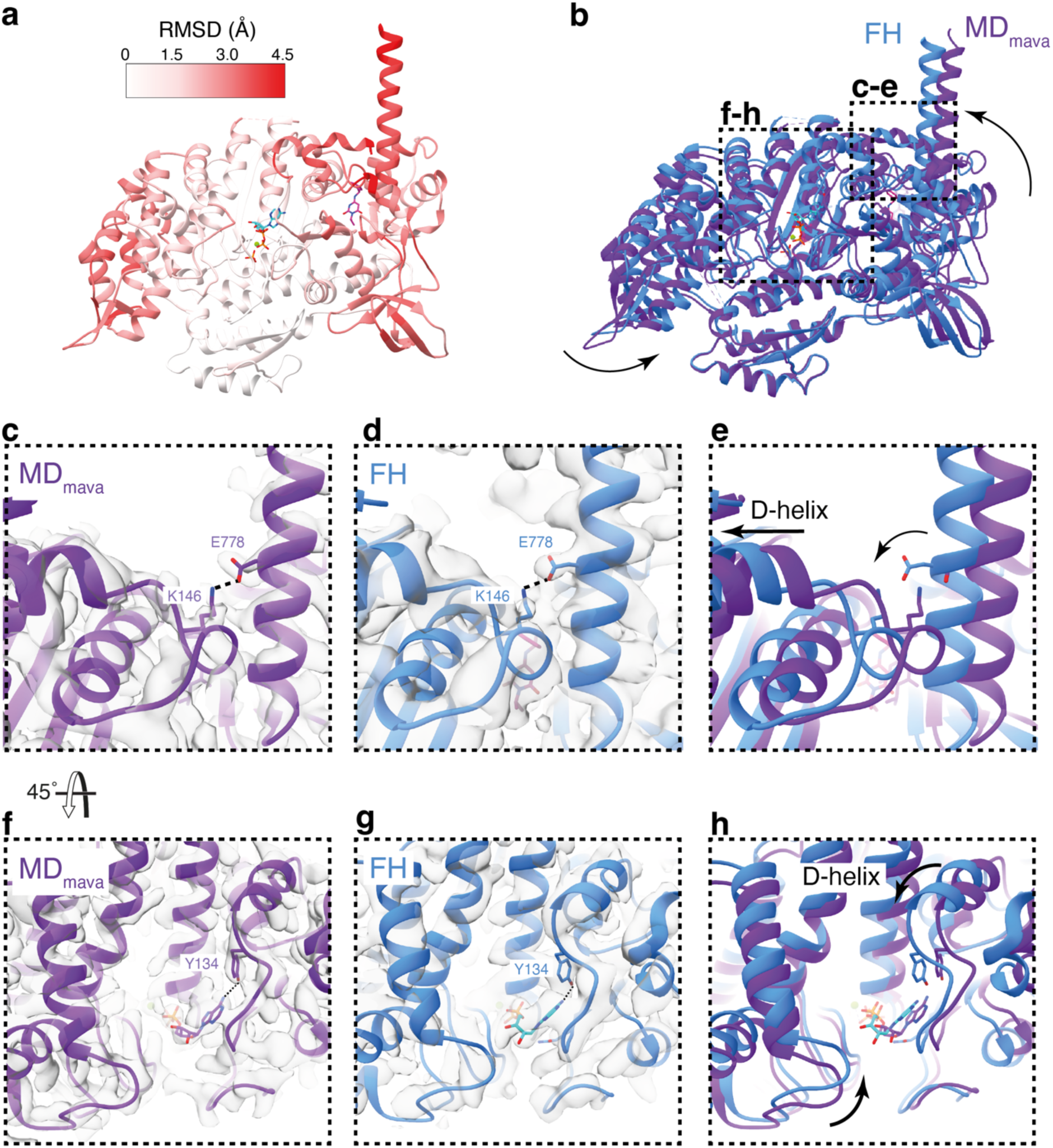
Mavacamten further inactivates the IHM free head. (a) RMSD comparison between MD_mava_ and IHM_mava_ FH aligned on the L50, coloured on IHM_mava_ FH model. (b) Overlay of MD_mava_ and IHM_mava_ FH blue highlighting domain movements. (c-e) Structural comparison of lever and D-helix conformation showing E778-K146 coupling hydrogen bond. (c) MD_mava_ model and cryoEM map (contour: 0.9), (d) IHM_mava_ FH model and cryoEM map (contour: 0.15) and (e) model overlay coloured as in (b). (f-H) Structural comparison of active site. (f) MD_mava_ model and cryoEM map (contour: 0.9), (g) IHM_mava_ FH model and cryoEM map (contour: 0.15) and (h) model overlay coloured as in (b).

Within the IHM_mava_ structure, the BH is further restrained by its interaction with S2 (Fig. 6). S2 predominantly interacts with the BH via the FH heavy chain directly strengthening the interaction between the two chains (Fig. 6a). The contact interfaces are predominantly charged interactions and can be split into three main regions on the BH motor; the OH loop, W-helix and HLH (Fig. 6b-f; Supplementary Movie. 3).

**Figure 6.**
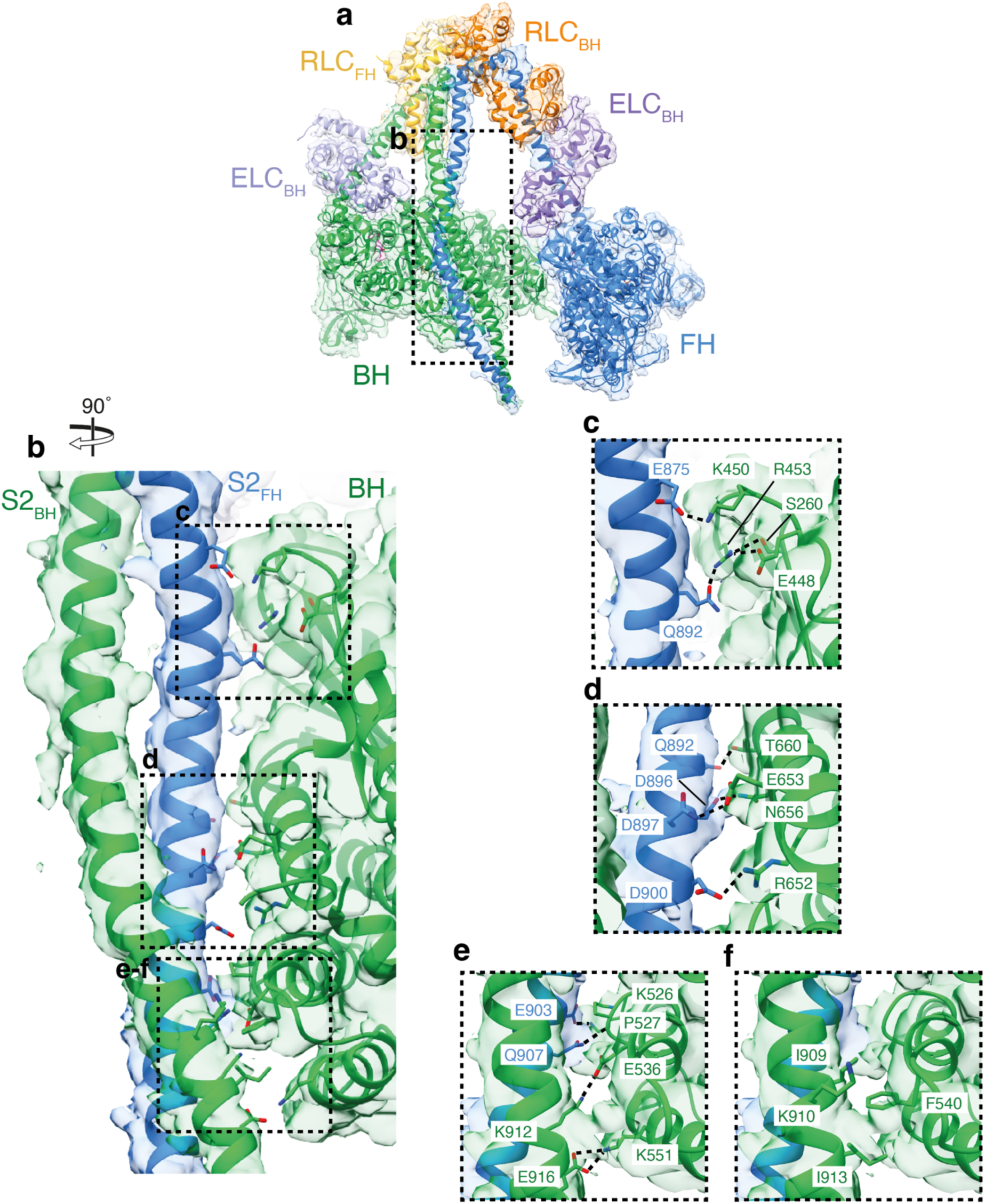
IHM_mava_ S2 novel interactions. (a) IHM_mava_ model in segmented cryoEM map coloured by chain (contour: 0.02): blocked head green, free head blue, blocked head ELC light purple, free head ELC purple, blocked head RLC yellow, free head RLC orange, mavacamten burgundy and the nucleotide in light blue. (b) Overview of IHM_mava_ S2-BH interactions (modelled utilising molecular dynamics in ISOLDE) in segmented cryoEM map (contour 0.01) BH green, FH blue. (c) S2-HO linker hydrogen bonding: E875_FH_-K450_BH_, Q892_FH_-R453_BH_ and R453_BH_-E448_BH_. (d) S2-W helix hydrogen bonding: Q892_FH_-T660_BH_, D896_FH_-N656_BH_, D897_FH_-N653_BH_ and D900_FH_-R652_BH_. (e-f) S2-HLH interactions (e) hydrogen bonding: E903_FH_-P527_BH_ backbone, Q907_FH_-K526_BH_, K912_BH_-E536_BH_ and E916_BH_-K551_BH_ (f) hydrophobic interactions: I909_BH_, I913_BH_ F450_BH_, and the aliphatic side chain backbone of K910_BH_.

The S2 in our IHM_mava_ structure adopts a curved conformation more closely resembling that observed in the thick filament structures^13,14,31^ over the previous single particle report of IHM^40^ (Extended Data Fig. 10). This is unlikely to be an effect of mavacamten, but rather due to our use of a construct containing a native cHMM S2 domain sequence compared to that used to determine the prior IHM^40^ structure, which had a short S2 sequence, ending at K942, followed by a leucine zipper^40^.

Examination of the S2-BH interface in our IHM_mava_ structure shows that it contains many more HCM mutation sites (R435C,H,S,L^41–44^, R660N^45^, R652G^46^, K903K/G^47,48^, I909M^49^, I913K^50^) in addition to those previously reported in the IHM^40^ (K450E/T^51,52^, Q882E^53^, Q892K^54^). Thus, maintenance of this interface is likely crucial to regulate force production during cardiac contractions.

## Discussion

In this work we have shown three high resolution cryoEM structures, two cHMM motor domains with and without mavacamten bound, alongside the cHMM IHM in complex with mavacamten, containing the native substrate ADP.Pi. The open head cHMM motor domain structures in combination with our qXL-MS analysis and the new understanding of actin activation^18^, allow us to present a new mechanism through which mavacamten elicits its effect on the myosin mechanochemical cycle (Fig. 7). Primed motors show lever and cleft dynamics that allow them to undergo basal ATPase activity (Fig. 7a) and weakly associate with actin. This can be regulated by sequestration of myosin molecules into the IHM, which significantly reduces ATPase activity (Fig. 7b). Upon weak binding of myosin to actin (Fig. 7c), cleft closure, Pi release and powerstroke are accelerated, resulting in post-powerstroke actomyosin (Fig. 7d). Mavacamten stalls motor force generation by restraining lever position and inducing structural changes that stabilise ADP.Pi binding and limit cleft dynamics (Fig 7e). This in turn increases the abundance of the IHM by strengthening interactions at the motor-motor interface (Fig. 7f). We find no evidence to support the proposal that mavacamten destabilises the L50 to prohibit myosin weakly binding to actin^32^. Instead, in disordered relaxed motors that can weakly associate with actin (Fig. 7g), we propose that mavacamten stabilises ADP.Pi binding and reduces actin-binding cleft dynamics to limit cleft closure. This inhibits the weak-to-strong actin binding transition that is required for progression through the mechanochemical cycle (Fig. 7h).

**Figure 7.**
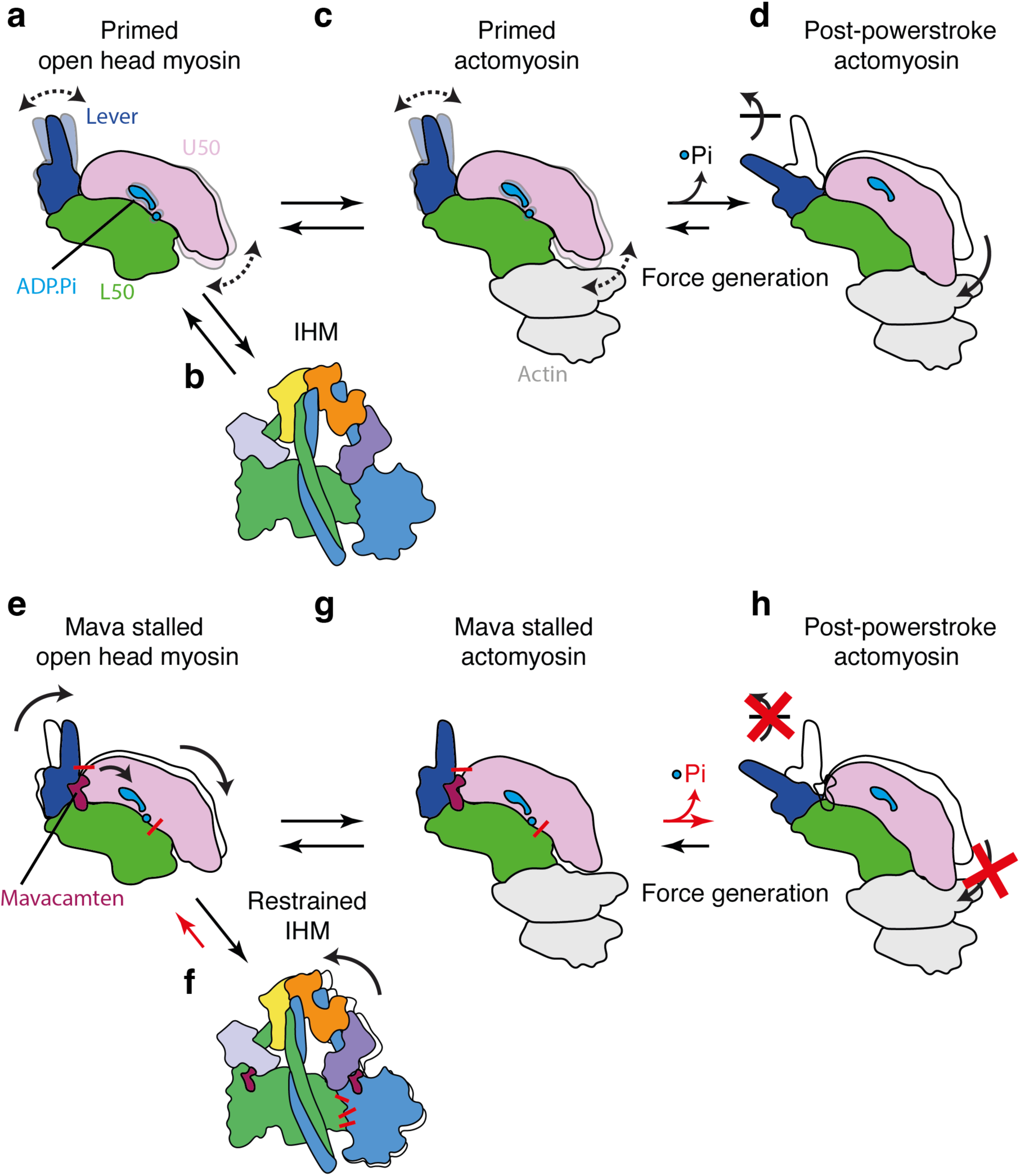
Overview of myosin force generation and mavacamten’s structural mechanism. (a-d) Schematic representation of βCM force generation: (a) Primed open head motor with both the lever and U50 dynamically exploring conformations able to transition to (b) the IHM preventing further force generation or (c) primed actomyosin. The primed actomyosin is initially weakly-bound but rapidly proceeds through the weak-to-strong actin binding transition, releasing phosphate and undergoing the force generating powerstroke resulting in formation of (d) post-powerstroke actomyosin. (e-h) Schematic representation of βCM force generation in the presence of mavacamten: (a) Mavacamten stalled open head motor with restrained lever and shifted U50 stabilising the Pi able to transition to (f) the restrained IHM, stabilised by increased motor-motor interactions. Mava stalled open head myosin may also interact with actin forming (g) stalled actomyosin. The stalled actomyosin does not readily undergo cleft closure (red-X) preventing phosphate release and its transition through lever swing (red-X) to (h) post-powerstroke actomyosin.

With deeper insight into the impact of mavacamten on motor mechanics, future work can begin to apply this in a clinically meaningful setting. The exploration of HCM mutations in the mavacamten binding site and key interfaces created by mavacamten will provide insight into the functional effects of the drug in patients with specific mutations. For example, the HCM mutations R712L and E774V would directly affect the mavacamten binding site likely rendering mavacamten less effective in patients with these mutations^55^. The HCM mutations R719W and R723G, within immediate proximity of the binding site, do mildly affect mavacamten binding^28,55^.

The newly formed network of motor-motor interactions in the presence of mavacamten also harbour several sites of pathogenic mutations: D382N^56^, K383V^57^, A393V^58^, R719W^59^, R719Q^60^, R719P^61^, Q734P^52^ and Q734E^62^. The presence of one of these mutations may lessen the effectiveness of mavacamten by reducing its IHM stabilising effects. Thus, further work is required to ascertain the effectiveness of mavacamten in the presence of disease-causing mutations within and surrounding the mavacamten binding site and at IHM contact interfaces. A deeper understanding of the mechanism through which mavacamten inhibits motor function in the context of disease may potentially allow us to predict patients who would be non-responders or who would potentially suffer unintended side effects from treatment with mavacamten, such as a significant reduction in systolic function. This would provide a path forward for personalised medicine alongside the development of more effective therapeutics.

## Methods

### Adenovirus manipulation

The human β-cardiac HMM (cHMM) used for cryoEM encodes residues 1-1137 of the *MYH7* gene (GenBank: AAA51837.1). This design includes 42 heptad repeats of the S2 domain with a FLAG tag added at the C-terminus (1138-1146) (Extended Data Fig. 2) and has been extensively analyzed^55,63^. A revised design was used for antibody capture for the motility assay that incorporates an epitope for an anti-S2 mAb (10F12.3)^64^. The antibody recognizes an epitope ‘AEKH RADLSRE’ spanning heptads 43 and 44 of MYH7. The epitope was added into coiled-coil S2 domain of β-cHMM followed by one additional heptad (45 heptads) and a FLAG tag at the C-terminus. The single-headed cS1 design encodes residues 1-834 MYH7 followed by a short linker fused to a GFP domain and a FLAG tag at the C-terminus. The expression cassette includes two IRES sequences for co-expression of human MYL3 (vLC1) and MYL2 (vLC2). The DNA sequences were assembled, inserted into an AdEasy shuttle vector, and sequenced (GeneWiz, Azenta Life Sciences, South Plainfield, NJ). Adenovirus plasmids were generated by recombination in E. coli strain BJ5183-Ad1 and the transgene inserts in the plasmids were sequenced. New viruses were packaged and amplified in Ad293 cells through five passages to produce high titer virus stocks^65^. The virus was harvested and purified by CsCl density gradient sedimentation yielding final virus titers of ∼10^11^ infectious units per mL (IU⋅mL^−1^) for infection of C2C12 cells and protein production.

### Muscle cell expression and purification of β-cardiac HMM and S1

Maintenance of the mouse myogenic cell line, C2C12 (CRL 1772; American Type Culture Collection, Rockville, MD), has been described in detail elsewhere^66^. Confluent C2C12 myoblasts were infected with replication defective recombinant adenovirus (AdcHMM2.0) at 2.7 X 10^8^ IU⋅mL^−1^ in fusion medium (89% DMEM, 10% horse serum, 1% FBS). Expression of recombinant myosin (cHMM or cS1) was monitored by accumulation of co-expressed GFP fluorescence in infected cells. Myocyte differentiation and GFP accumulation were monitored until the cells were harvested (198 – 264 hr). Cells were chilled, media removed, and the cell layer was rinsed with cold PBS. The cell layer was scraped into Triton extraction buffer: 100 mM NaCl, 0.5% Triton X-100, 10 mM Imidazole pH 7.0, 1 mM DTT, 5 mM MgATP, and protease inhibitor cocktail (Sigma-Aldrich, St. Louis). The cell suspension was collected in an ice-cold Dounce homogenizer and lysed with 15 strokes of the tight pestle. The cell debris in the whole cell lysate was pelleted by centrifugation at 17,000 x g for 15 min at 4°C. The Triton soluble extract was fractionated by ammonium sulfate precipitation using sequential steps of 0-30% saturation and 30-60% saturation. The myosin precipitates between 30-60% saturation of ammonium sulfate. The recovered pellet was dissolved in and dialyzed against 50 mM Tris, 150 mM NaCl, pH 7.4, 0.5 mM MgATP for affinity purification of the FLAG-tagged myosin on M2 mAb-Sepharose beads (Sigma-Aldrich). Bound myosin was eluted with 0.1 mg⋅mL^−1^ FLAG peptide (Sigma-Aldrich). Protein was concentrated and buffer exchanged on Amicon Ultracel-10K centrifugal filters (Millipore; Darmstadt, Germany), dialyzed exhaustively into 10 mM MOPS, 100 mM KCl, 1 mM DTT before a final centrifugation at 300,000 x g for 10 min at 4°C. Aliquots were drop frozen in liquid nitrogen and stored in vapor phase at – 147°C. The sequence of the β-cHMM and cS1 preparations used in this study were confirmed by LC-MS/MS of protein digests. Bound light chains are those that are constitutively expressed in the C2C12 cells (MLC1/MLC3 and rLC2) or co-expressed with the cS1gfp (vLC1 and vLC2).

### In vitro motility assay

Measurement of in vitro motility of the S2 epitope tagged cHMM was done as previously described for skeletal muscle myosin^64,67^. Nitrocellulose-coated glass coverslips were incubated with 0.1 mg/mL of the anti-S2 mAb, 10F12.3, in PBS followed by blocking the surface with 1% BSA/PBS. The cS1gfp was tethered with the anti-GFP mAb 3E6 (Invitrogen, ThermoFisher) bound to nitrocellulose-coated glass coverslips. The β-cHMM and cS1 proteins were diluted to 20 µg/mL in motility buffer (MB) (25 mM imidazole, pH 7.8, 25 mM KCl, 4 mM MgCl_2_, 1 mM MgATP, 1 mM DTT) supplemented with 1% BSA (MB/BSA). The antibody-coated coverslips were incubated with the proteins for 2 hr in a humidified chamber at 4 °C. The coverslips were washed sequentially with MB/BSA and 3-times with MB before blocking with 0.5 μM unlabelled F-actin (5 min) and two final washes with MB. The coverslips were mounted on a 15-μL drop of 2 nM rhodamine-phalloidin–labelled actin in a modified motility buffer (with 7.6 mM MgATP, 50 mM DTT, 0.5% methyl cellulose, 0.1 mg/mL glucose oxidase, 0.018 mg/mL catalase, 2.3 mg/mL glucose) in a small parafilm ring fixed on an alumina slide with vacuum grease. The chamber is observed with a temperature-controlled stage and objective set at 32 °C on an upright microscope with an image-intensified CCD camera capturing images to an acquisition computer at 5– 30 fps depending on assay parameters. Movement of actin filaments in 2–3 fields/slide for 500 frames/field of continuous imaging were captured and analyzed with semiautomated filament tracking programs as previously described^67^. The trajectory of every filament with a lifetime of at least 10 frames is determined; the instantaneous velocity of the filament moving along the trajectory, the filament length, the distance of continuous motion and the duration of pauses are tabulated. A weighted probability of the actin filament velocity for hundreds of events is fit to a Gaussian distribution and reported as a mean velocity and SD for each experimental condition.

### Negative stain head counting analysis

cHMM was first prepared by crosslinking with bis(sulfosuccinimidyl)suberate (BS3) for 30 minutes at 25 °C in the following conditions: 2 µM cHMM, 50 µM mavacamten (5 % DMSO), 1 mM BS3, 1 mM EGTA, 1 mM ATP, 2 mM MgCl_2_, 56 mM KCl, 10 mM MOPS pH 7.2. The reaction was then quenched with a 100 mM final concentration of Tris pH 8 preventing further crosslinking. Crosslinking was confirmed by SDS-PAGE analysis (Supplementary Fig. 3).

Negative stain grids were prepared using the flicking method^68^ by applying 5 µl of crosslinked cHMM diluted 5-fold in buffer to a negative stain grid (produced in-house) glow discharged for 30s (PELCO easiGlow™) prior to use. The excess HMM was then flicked off and a drop of 2 % uranyl acetate applied. The excess was again flicked off and the addition of a 2 % uranyl acetate drop repeated four times. Finally, the grid was blotted with Whatman no. 42 Ashless filter paper and dried. Negative stain image collection was performed using the FEI Tecan F20 equipped with a FEI CETA (CMOS CCD). Images were collected at 29,000x magnification at a defocus of −1 to −3 µm. Images were double blinded prior to counting of open and IHM-like particles. Images were only included in the analysis if a minimum of 250 particles were detected within the image such that the sample size was sufficient to obtain 80 % power with a 95 % confidence interval. Data was then plotted into box plots and the significance in change between populations was calculated using a two-tailed student t-test in GraphPad Prism.

### CryoEM grid preparation and data collection

Prior to cryoEM grid preparation cHMM was crosslinked with BS3 using the same protocol as for negative stain with altered buffer conditions: 2 µM cHMM, 50 µM mavacamten (5 % DMSO), 1 mM BS3, 1 mM EGTA, 1 mM ATP, 2 mM MgCl_2_, 50 mM KCl, 10 mM MOPS pH 7. Grids were prepared using the Vitrobot Mark IV (Thermo Fisher). 3µl of the BS3 crosslinked cHMM diluted 2-fold in buffer was applied to an UltrAuFoil^TM^R 1.2/1.3 300 mesh gold grid (Quantifoil) for motor domain reconstruction and a UC-A on Lacey 400 mesh Cu continuous carbon grid (Agar scientific) for IHM reconstruction, glow discharged for 30 seconds at 10 mA prior to use (PELCO easiGlow™). Grids were then blotted with Whatman no. 42 Ashless filter paper (Agar Scientific, UK) for 1s with a force of 6, and wait time of 2s at 8 °C and 95 % humidity before vitrification in liquid ethane. Data was collected on a FEI Titan Krios (Astbury Biostructure Laboratory, University of Leeds) operating at 300 kV equipped with a Falcon 4 direct electron camera with a specimen pixel size of 0.82 Å. Micrographs were collected using EPU acquisition software at 96,000x magnification with a total dose of 43e^−^/Å^2^ and a defocus range of −1.2 to −3.0 µm. Total micrographs for each reconstruction were: MD 9,948 over one session, MD_mava_ 21,395 over two sessions and IHM_mava_ 34,287 over three sessions.

### CryoEM data processing and model building

MD and MD_mava_ single motor domain image processing was carried out in RELION 4.0^69^ with subsequent processing in cryoSPARC V4.2^70^. Raw movies were imported into RELION for motion correction using MotionCor2^71^ and CTF estimation using CTFFIND-4.1^72^. Automated particle picking was then performed using Topaz initially implementing the general model then a trained Topaz^73^ model on selected single motor domain 2D classes, this was repeated for each dataset. Particles were extracted in a box size of 360^2^ pixels centred on individual motor domains. The resulting particles from both collections were then combined and classified using cryosparc reference-free 2D classification^70^. Classes resembling single motor domains were selected for further classification. An initial model was generated using cryoSPARC’s ab-initio reconstruction into five classes. The resulting maps were then refined through heterogeneous, homogeneous and finally non-uniform refinement^70,74^ on combined selected classes with a final particle stack of 88,809 MD and 200,487 MD_mava_. The resulting map was then sharpened using a negative B-factor automatically determined by cryoSPARC and local resolution estimation was calculated in cryoSPARC^70^. IHM_mava_ image processing followed the same pipeline except IHM 2D classes were selected to train a topaz^73^ model and only three ab-initio^70^ classes were used for initial model generation. A total of 197,869 particles contributed to the final map.

To interpret the cryoEM maps, atomic models for the single motor domain and IHM were produced. Homology models for human βCM heavy and light chains were generated using the smooth muscle myosin shutdown structure (PDBID: 67Z4)^75^. These models were then truncated at residue 796 (within the LCD) and flexibly fit to the single motor domain density using ISOLDE^34^. Phenix real space refinement was then performed followed by adjustments in Coot^76^ and ISOLDE^34^, this was repeated until satisfied. The IHM model was then built using two MD_mava_ motor domain models joined to the homology modelled LCD, S2 and light chains. The model was flexibly fitted into the IHM map by use of ISOLDE and refined using Phenix real space refinement with final adjustments in Coot^76^ and ISOLDE^34^.

### qXL-MS sample preparation, measurement, data preparation and analysis

Purified cS1 (25 µL, final concentration of 4 µM in 10 mM MOPS, pH 7.3, 50 mM KCl, 1 mM MgCl_2_, 0.34 mM DTT, 1 mM EGTA, 1 mM ATP) was incubated with 50 mM of Mavacamten or DMSO control for 30 minutes at 25 °C. DSBU (600 µM; 149 molar fold excess) was added, or DMSO control, and was allowed to react for 20 minutes at 25 °C (final DMSO concentration of 1.6 % v/v). The reaction was quenched by the addition of Tris (1 M, pH 7.3) to final concentration of 20 mM and incubation at room temperature for 15 minutes. Samples were flash frozen for storage prior to digestion. Crosslinking was confirmed by SDS-PAGE analysis (Supplementary Fig. 3)

Three replicates of both crosslinked and non-crosslinked control samples (≈14.5 µg) were processed for MS analysis using S-Trap micro spin columns (Protifi) as described previously^77^ after which the peptides were resuspended in 5 % v/v acetonitrile/0.1 % v/v formic acid. Samples, ≈10 % of the final volume of each replicate, were analyzed on an Vanquish Neo LC (Thermo) coupled to an Orbitrap Eclipse mass spectrometer (Thermo) in positive ion and DDA mode. Separation of peptides was performed using PepMap Neo C18 trap cartridge (Thermo Fisher Scientific, 174500) before using the EASY spray C18 column (Thermo Fisher Scientific, ES903). Elution of peptides from the column was achieved using a gradient elution of a 7.5-42.5 % (v/v) solvent B (0.1 % (v/v) formic acid in acetonitrile) in solvent A (0.1 % (v/v) formic acid in water) over 97.5 min at 250 nl min^−1^. The eluate was infused into the instrument using an EASY-Spray nanoelectrospray ionization source.

An online exclusion list was generated from the MS1 measurement of a non-crosslinked control sample using the AcquireX AB workflow editor (Thermo Fisher Scientific application module) and was applied when performing MS analysis of the crosslinked samples. The Full Scan and Deepscan methods were adapted from previously reported method^77^ where the ‘Full scan’ method that lacks an MS^2^ product ion scan found in the Deepscan method both were Xcalibur AcquireX AB enabled and identical chromatographic parameters.

The .RAW MS files produced were processed directly in FragPipe (v21.1)^78^, without conversion to mzML, subject to a mass offset search using MSFragger (v4.0)^78^ where default “Mass-Offset-Common-PTMs” workflow was loaded and amended. False discovery rate (FDR) at the protein/peptide/ion level was set to 1 % and tolerances for precursors/fragments set to 10 ppm. The generated calibrated MZML files were then taken forward as the generated interact.pep.xml file.

MeroX (v2.0.1.4) was used to search the replicate files individually for crosslinks and monolinks (from residues K,S,T and Y) against FASTA files for the S1GFP construct and associated LCs, as used for the MSfragger search, without the appended decoys and common contaminants from MSfragger, with a 1.0 % FDR and 50 score cutoff. Precursor and fragment ion precision was set to 10 ppm with a 2.0 signal to noise ratio. A single reporter ion fragment was allowed to be missing in the database search. Dead-end crosslinks were allowed to react with Tris as well as allowing intrapeptide crosslinks and the neutral loss of C_4_H_7_NO.

The software package Skyline^79^ (MacCoss Lab Software, v23.1.0.455) was used to quantify the crosslinks and monolinks found by MeroX using a modified process laid out by Chen and Rappsiliber^37^ and since improved by Sinz^80^ and separately by Jaing et al^81^. The six replicate MeroX search result .ZHRM files results were converted to the ProXL XML files (https://github.com/yeastrc/proxl-import-merox).

All crosslinks and monolink peptides were manually curated and all peaks of all transitions were manually inspected and aligned in Skyline. Crosslinks and monolinks were kept for quantitation if at least two of the replicates had convincing MS2 assignment in MeroX (spectral matched scored over 125 and near complete MS2 peak identification), and all three of the DMSO or Mavacamten replicates had well defined MS1 peaks with matching retention times and strong Skyline MS2 signal. Further to this, cross- or mono-links with peptides where DSBU is potentially labelled on different residues but the assignment or resolution of the peaks poor were determined by cross-referencing the MeroX annotated MS2 spectrum with the Skyline MS2 transitions. In some cases, where the baseline separation of the peaks was good but where no clear determination could be made, the highest scoring option within MeroX (annotated MS2 spectrum) was used and all MS1 signals were assigned to this assignment.

The signal intensity for each peptide for each replicate was normalised, as described by Chen and Rappsiliber^37^, using the total non-DSBU reacted signal intensity of each replicate. The interact.pep.xml file from the MSfragger search was used to import the search of the six replicates into a separate Skyline file. Monolink signal intensity was combined from peptides that contain the same monolinked residue regardless of peptide charge state or length. Similarly, crosslink signal intensity was combined from dipeptides that bear the same residue to residue crosslink.

An aligned interpeptide crosslink, intrapeptide crosslink or monolink was determined to have changed significantly in the presence or absence of Mavacamten by conducting pairwise comparisons via a single tailed, homoscedastic t-test using relative signal intensity (Supplementary Material 2). The significance was measured by protein fold changes >2 and p < 0.05.

### Data availability

The electron density maps and atomic models for MD, MD_mava_ and IHM_mava_ have been deposited into EMDB, with accession codes EMD-51721, EMD-51720 and EMD-51719, and the PDB files with accession codes 9GZ3, 9GZ2 and 9GZ1, respectively. The XL-MS dataset generated in this study, including experimental settings and XL identification results, has been deposited to the ProteomeXchange Consortium via the PRIDE^82^ partner repository with the dataset identifier PXD059316.

The following models were used for comparison purposes in our study, Bovine S1 cardiac myosin in complex with mavacamten PDB ID: 8QYQ and the folded-back state (IHM), PDB ID: 8ACT. In addition to the above models, the following cryoEM maps were used for comparisons: the folded-back state (IHM) EMDB ID: 15353, the human cardiac thick filament EMDB ID: 29726 and the human cardiac thick filament EMDB ID: 40471.

## Acknowledgements

We would like to thank Prof. Peter Knight, Prof. Michelle Peckham, and Prof. Stephen Muench for their valuable insight and comments on the manuscript. We thank the members of the cryoEM and spectrometry community at Leeds for their help and guidance. All EM data was collected at the Astbury Biostructure facility funded by the University of Leeds (UoL ABSL award) and Wellcome (108466/Z/15/Z and 221524/Z/20/Z). All mass spectrometry data was collected at the Biomolecular Mass Spectrometry Facility at the University of Leeds with funding from Wellcome (223810/Z/21/Z) for the Eclipse mass-spectrometer and technical support from S. R. Ganji. This work was supported by a British Heart Foundation Jacqueline Murray Coomber Fellowship (FS/0/21/34704) and Royal Society research grant (RGS\R1\231276) awarded to C.A.S. and a NIH grant (5R01HL157997) to D.A.W., S.N.M is supported by a School of Molecular and Cellular Biology, University of Leeds, funded PhD studentship.

## Author contributions

C.A.S designed the project. B.B and D.A.W produced the beta cardiac myosin. D.A.W performed in vitro motility assay and data analysis. S.N.M performed the cryoEM grid preparation, screening, optimisation and data collection. S.N.M performed the cryoEM data processing, model building and validation. S.N.M and C.A.S performed cryoEM data analysis and interpretation. J.R.T.P performed crosslinking spectrometry preparation, data collection and data processing. J.R.T.P and C.A.S performed crosslinking spectrometry data interpretation. S.N.M performed main figure, supplementary figure and movie generation. S.N.M, J.R.T.P and C.A.S wrote the manuscript. All authors discussed the results and commented on the manuscript.

---

**Supplementary Table 1.** Inhibition of cardiac myosin motor activity by mavacamten.

| cHMM | cS1 |
| --- | --- |
| IC<sub>50</sub> = 0.14 ± 0.01 μM | IC<sub>50</sub> = 0.62 ± 0.07 μM |
| V<sub>o</sub> = 1.26 ± 0.04 μm/s | V<sub>o</sub> = 2.05 ± 0.05 μm/s |
| V<sub>i</sub> = 1.12 ± 0.04 μm/s | V<sub>i</sub> = 1.77 ± 0.07 μm/s |
| V<sub>min</sub> = 0.14 μm/s | V<sub>min</sub> = 0.28 μm/s |

**Supplementary Table 2:**
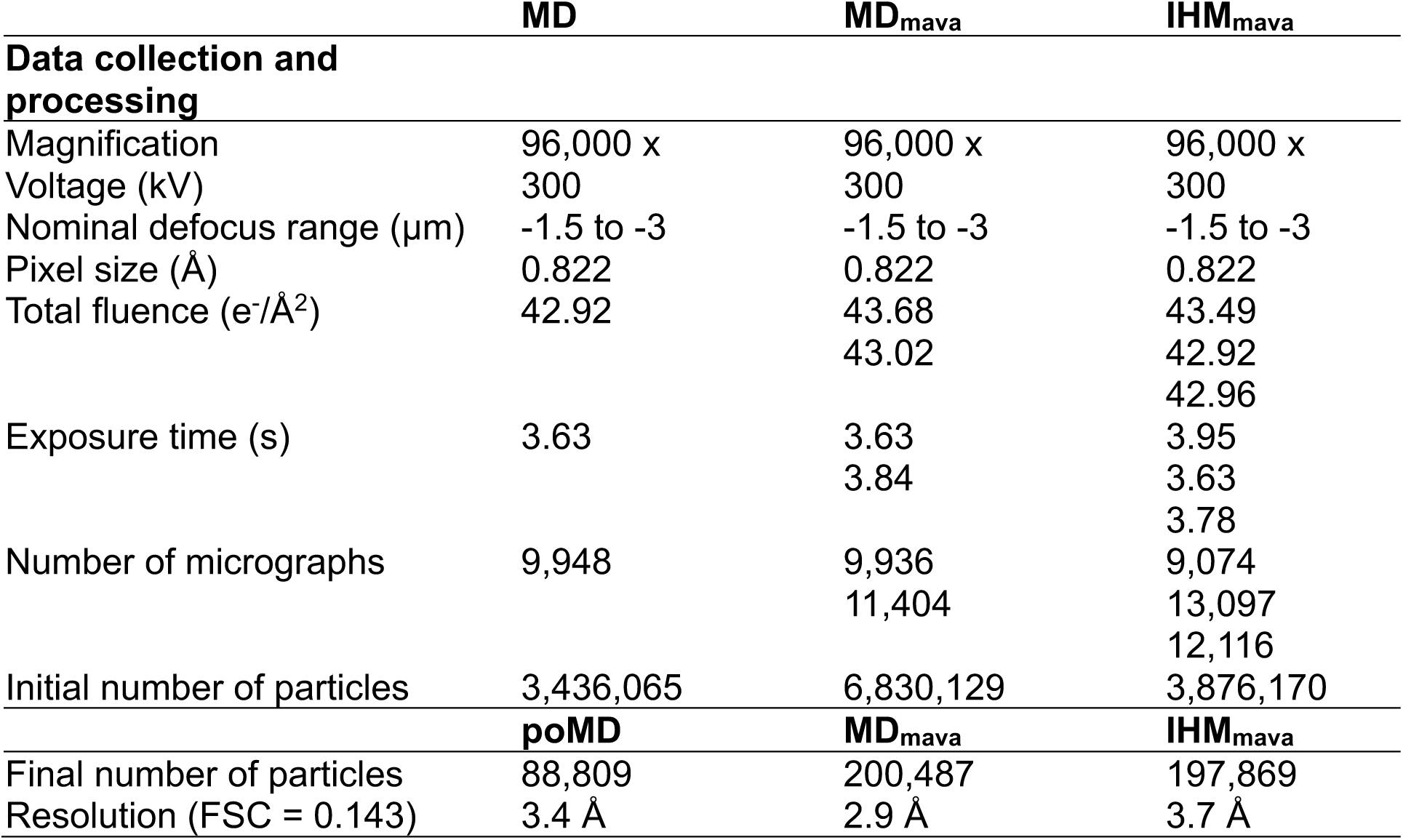
Data collection and processing statistics for MD, MD_mava_ and IHM_mava_ EM structures. Where multiple collections were combined individual values for each collection are listed.

|  | MD | MD<sub>mava</sub> | IHM<sub>mava</sub> |
| --- | --- | --- | --- |
| **Data collection and processing** |  |  |  |
| Magnification | 96,000 x | 96,000 x | 96,000 x |
| Voltage (kV) | 300 | 300 | 300 |
| Nominal defocus range (μm) | -1.5 to -3 | -1.5 to -3 | -1.5 to -3 |
| Pixel size (Å) | 0.822 | 0.822 | 0.822 |
| Total fluence (e<sup>-</sup>/Å<sup>2</sup>) | 42.92 | 43.68 | 43.49 |
|  |  | 43.02 | 42.92 |
|  |  |  | 42.96 |
| Exposure time (s) | 3.63 | 3.63 | 3.95 |
|  |  | 3.84 | 3.63 |
|  |  |  | 3.78 |
| Number of micrographs | 9,948 | 9,936 | 9,074 |
|  |  | 11,404 | 13,097 |
|  |  |  | 12,116 |
| Initial number of particles | 3,436,065 | 6,830,129 | 3,876,170 |
|  | **poMD** | **MD<sub>mava</sub>** | **IHM<sub>mava</sub>** |
| Final number of particles | 88,809 | 200,487 | 197,869 |
| Resolution (FSC = 0.143) | 3.4 Å | 2.9 Å | 3.7 Å |

**Supplementary Table 3:**
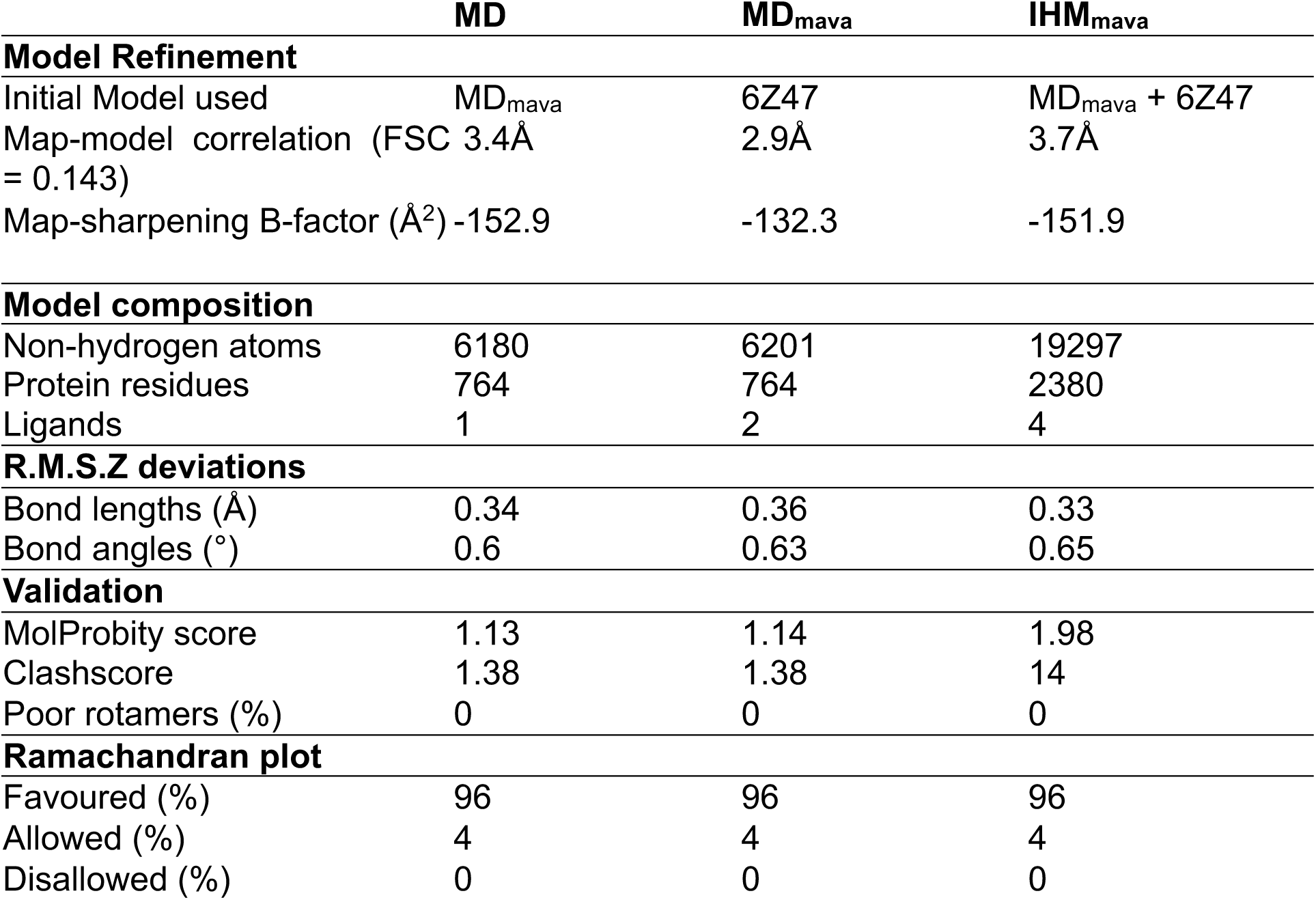
Model building and refinement statistics for MD, MD_mava_ and IHM_mava_ EM data.

**Supplementary Table 4:** Interdomain crosslinks annotated on Fig. 4. C⍺-C⍺ distances that change >2 Å between models are highlighted in green. Positive to negative log2fold change in crosslink intensity on addition of mavacamten is indicated on a colour scale from red to blue.

| Protein1 | Domain 1 | Residue 1 | Protein 2 | Domain2 | Residue 2 | Cα-Cα distance Å MD (BHapo) | Cα-Cα distance Å MDmava (BHmava) | Log2 fold change in intensity + mavacamten | T-test | Indication with mava | Figure Panel |
| --- | --- | --- | --- | --- | --- | --- | --- | --- | --- | --- | --- |
| MYH7 | U50 | S205 | ELC | EF-hand 2 | T147 | 18.2 | 15.3 | 2.65 | 0.004 | increased proximity | a/c/d |
| MYH7 | U50 | K207 | ELC* | EF-hand 2 | K142 | 18.3 | 15.5 | 1.69 | 0.005 | increased proximity | c/d |
| MYH7 | U50 | T255 | ELC* | EF-hand 2 | K142 | 12.2 | 7.9 | 1.50 | 0.026 | increased proximity | a/c/d |
| MYH7 | U50 | K450 | ELC | EF-hand 2 | K142 | 26.9 | 22.8 | -1.19 | 0.016 | decreased reactivity | b/c |
| MYH7 | U50 | S205 | RLC | EF-hand 2 | K115 | 47.8 (29.7) | 42.5 (22.4) | 5.96 | 0.004 | increased stability | a/c/d |
| MYH7 | U50 | S205 | RLC | loop region | K111 | 42.4 (25.5) | 37.9 (18.8) | 3.02 | 0.047 | increased stability | a/c/d |
| MYH7 | LCD | K803 | RLC | EF-hand 2 | K111 | 21.5 | 21.5 | 3.02 | 0.002 | increased stability | a/c/d |
| MYH7 | LCD | K803 | RLC | EF-hand 2 | K115 | 21.4 | 21.4 | 1.83 | 0.001 | increased stability | a/c/d |
| MYH7 | LCD | K837 | RLC | - | K165 | 11.9 | 11.9 | 3.81 | 0.006 | increased stability | a |
| MYH7 | LCD | K825 | RLC | - | K165 | 19.2 | 19.2 | 2.91 | 0.024 | increased stability | a |
| MYH7 | LCD | K835 | RLC | - | K160 | 12.2 | 12.2 | 2.79 | 0.008 | increased stability | a |
| MYH7 | LCD | K835 | RLC | - | K165 | 9.5 | 9.5 | 2.19 | 0.008 | increased stability | a |
| MYH7 | LCD | K835 | RLC | EF-hand 1 | K62 | 18.4 | 18.4 | 1.51 | 0.012 | increased stability | a |
| MYH7 | NTD | K34 | ELC | loop region | K98 | 46 | 47.7 | -2.43 | 0.000 | reduced dynamics | b/e |
| MYH7 | NTD | K21 | ELC | loop region | K98 | 28.3 | 27.9 | -2.45 | 0.001 | reduced dynamics | b/e |
| MYH7 | NTD | K50 | ELC | loop region | K98 | 49.9 | 51 | -4.19 | 0.006 | reduced dynamics | b/e |
| MYH7 | NTD | K34 | MYH7 | U50 | K83 | 15.9 | 15.7 | 3.90 | 0.000 | increased stability | a/e |
| MYH7 | NTD | K58 | MYH7 | L50 | K565 | 39.9 | 40.2 | 2.83 | 0.000 | increased reactivity | a/e |
| MYH7 | NTD | K50 | MYH7 | L50 | K707 | 29.9 | 28.8 | -0.72 | 0.020 | reduced reactivity | b/e |
| MYH7 | L50 | K707 | ELC | EF-hand 1 | K72 | 60.6 | 60.4 | -1.61 | 0.011 | reduced dynamics | b/e |
| MYH7 | L50 | K707 | MYH7 | CON | Y756 | 12.4 | 12.3 | 6.20 | 0.013 | increased stability | a/e |
| MYH7 | L50 | K707 | MYH7 | CON | K761 | 20.8 | 20.9 | 3.60 | 0.019 | increased stability | a/e |
| MYH7 | L50 | K707 | MYH7 | CON | K757 | 13.1 | 13.1 | 2.64 | 0.002 | increased stability | a/e |
| MYH7 | U50 | K405 | MYH7 | L50 | T646 | 20.2 | 20.2 | 2.74 | 0.001 | increased stability | a/e |
| MYH7 | U50 | K413 | MYH7 | L50 | K598 | 19.4 | 19.3 | 1.78 | 0.001 | increased stability | a/f |
| MYH7 | U50 | K405 | MYH7 | L50 | K598 | 18.1 | 17.6 | 1.40 | 0.004 | increased stability | a/f |
| MYH7 | U50 | K450 | MYH7 | L50 | T646 | 34.9 | 34.7 | -2.08 | 0.005 | reduced dynamics | b/f |
| MYH7 | U50 | T449 | MYH7 | L50 | K707 | 45.6 | 45.2 | -2.21 | 0.004 | reduced dynamics | b/f |
| MYH7 | U50 | K450 | MYH7 | L50 | K657 | 24.5 | 24.8 | -3.95 | 0.003 | reduced reactivity | b/f |

## Extended data figures

**Extended Data Fig. 1.**
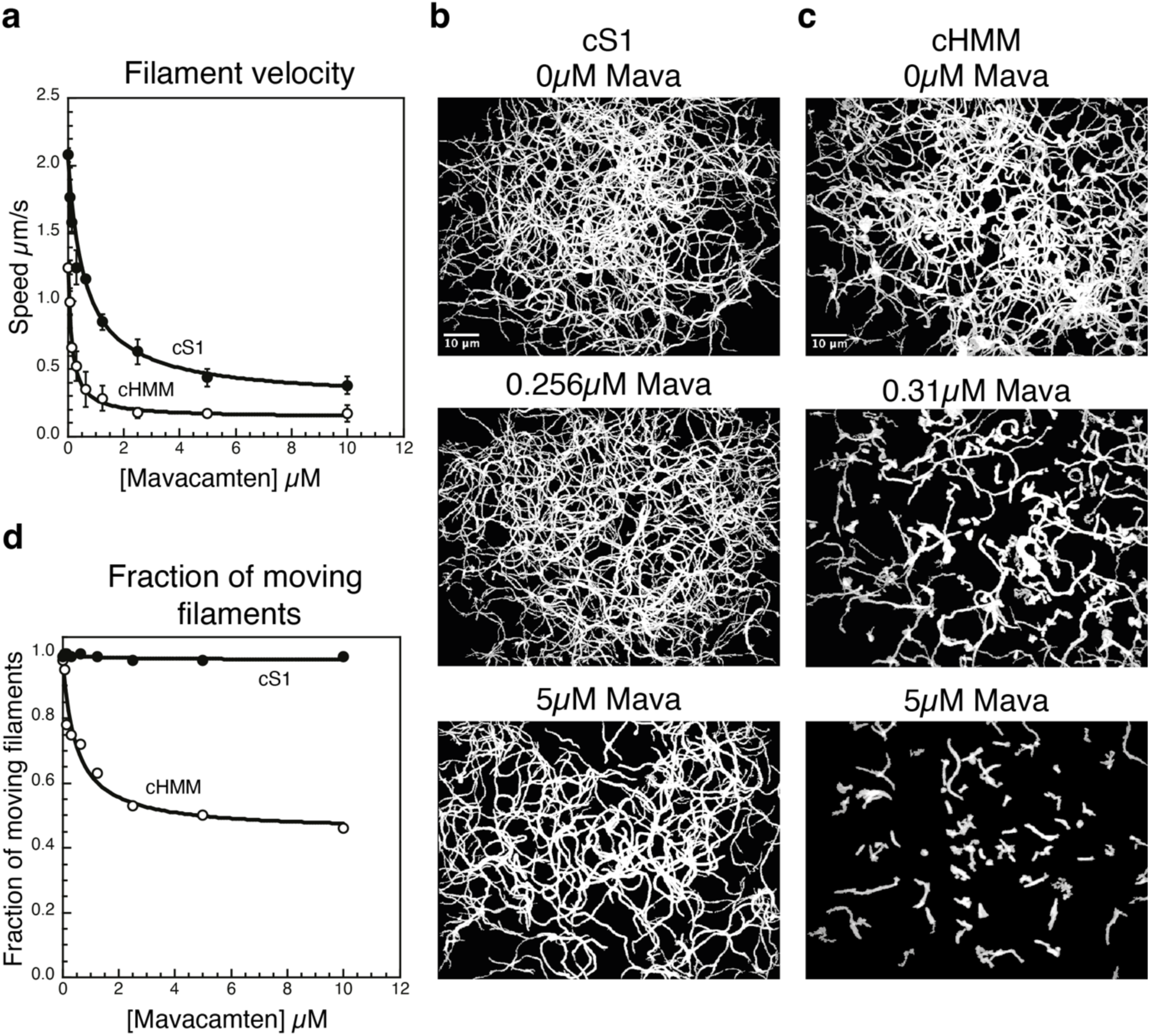
Mavacamten inhibits the gliding velocity of actin filaments more effectively for cHMM compared to cS1. (a) Actin filament gliding velocity for cS1 and cHMM over a titration of mavacamten. The filament speed powered by cS1 is slowed up to 86 % by mavacamten with an IC_50_ of 0.62 µM. Comparatively, the filament speed powered by cHMM is slowed by 90 % with a 4-fold lower IC_50_ of 0.14 µM. (b-c) Summed plot of actin filament movement over 100 seconds of motility for (b) cS1 and (c) cHMM at mavacamten concentrations of 0 µM, ∼0.3 µM and 5 µM. (d) Analysis of the number of moving filaments for cS1 and cHMM over a titration of mavacamten. Mavacamten has no impact on the fraction of filaments moving for cS1 however, mavacamten decreases the fraction of moving filaments for cHMM in a concentration dependent manner.

**Extended Data Figure 2.**
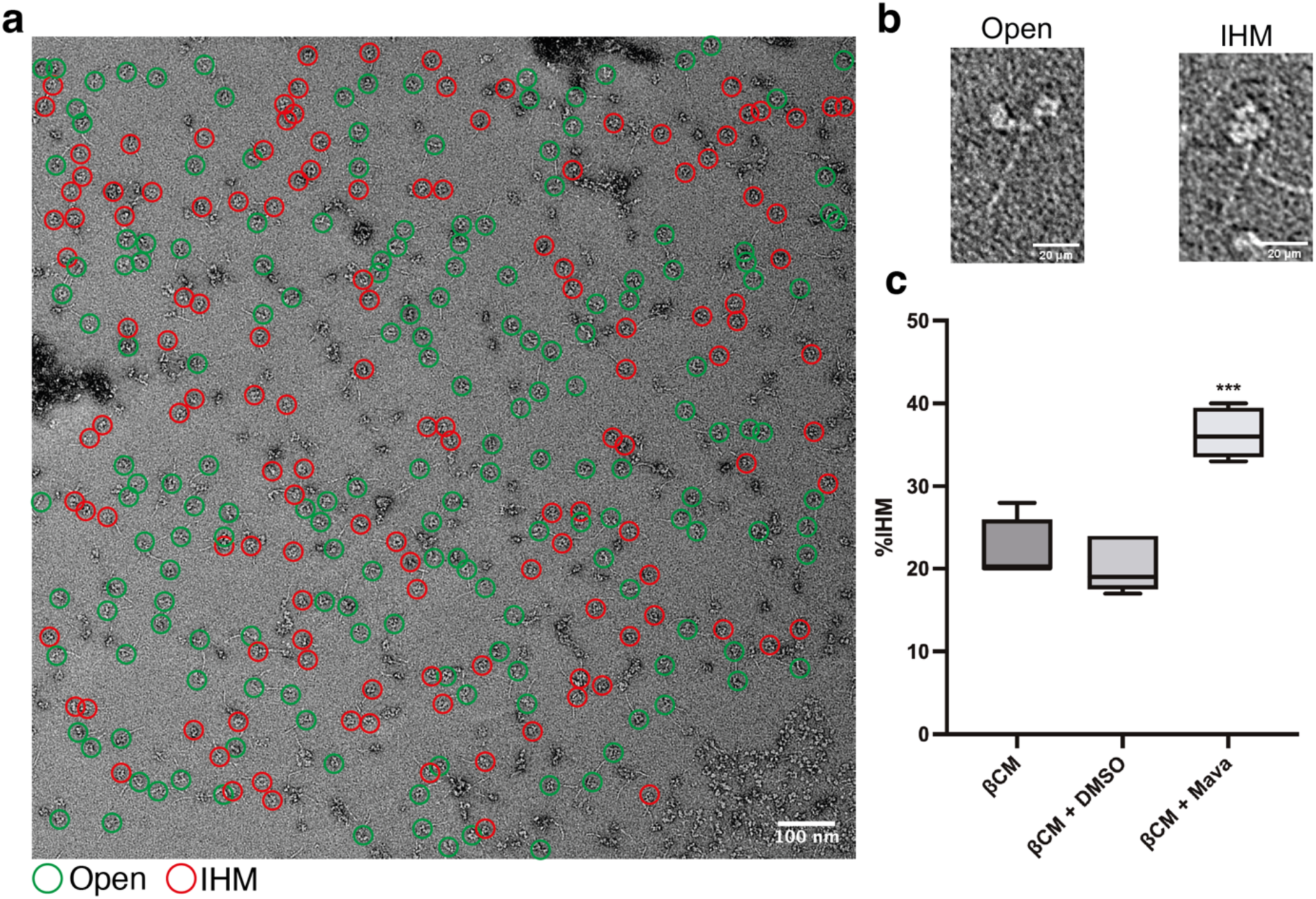
Direct observation of IHM stabilisation due to mavacamten. (A) Representative negative-stain EM micrograph used during head counting assay. Counted ßCM particles highlighted by coloured circle, green: open, red: IHM. (b) Representative negative stain open and IHM ßCM molecule. (c) Box plot showing percentage of IHM over 5 biological replicates. The ßCM control resulted in a median of 20 % with a first and 3rd quartile of 20 % and 24 % respectively. ßCM containing 2.5 % v/v DMSO showed a median of 19 % with a first and third quartile of 18 % and 24 % with no significant difference to the ßCM control determined by an unpaired two-tailed student t-test. Mavacamten shows a median of 36 % with a first and third quartile of 34 % and 39 %, a significant increase in % IHM formation with a P-value of = 0.0002 determined by an unpaired two-tailed student t-test with respect to the ßCM control.

**Extended Data Figure 3.**
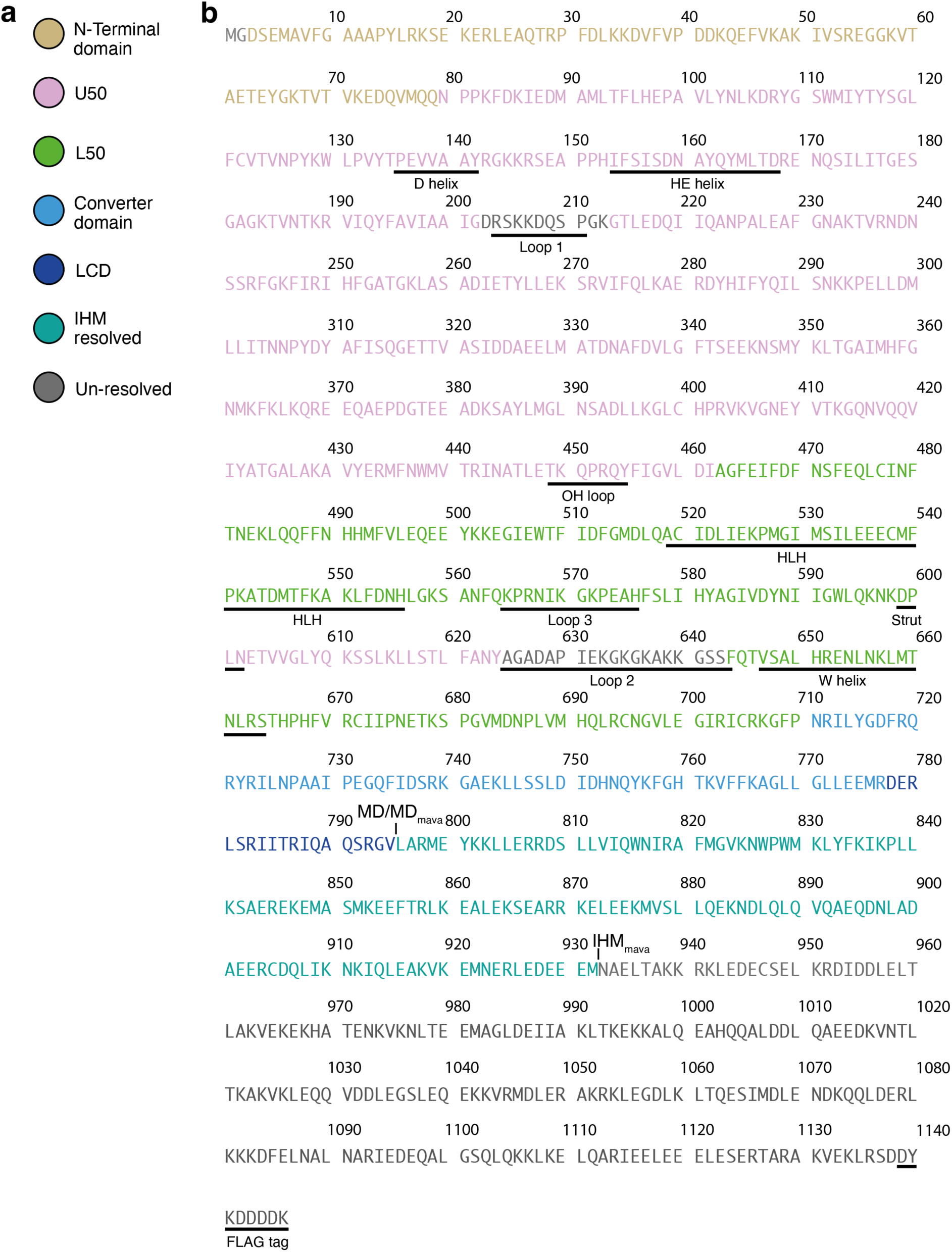
cHMM heavy chain sequence and sub domains. (a) Key for cHMM sequence (b) cHMM heavy chain sequence highlighting resolved sub-domains and key structural regions with C-terminal flag tag.

**Extended Data Figure 4.**
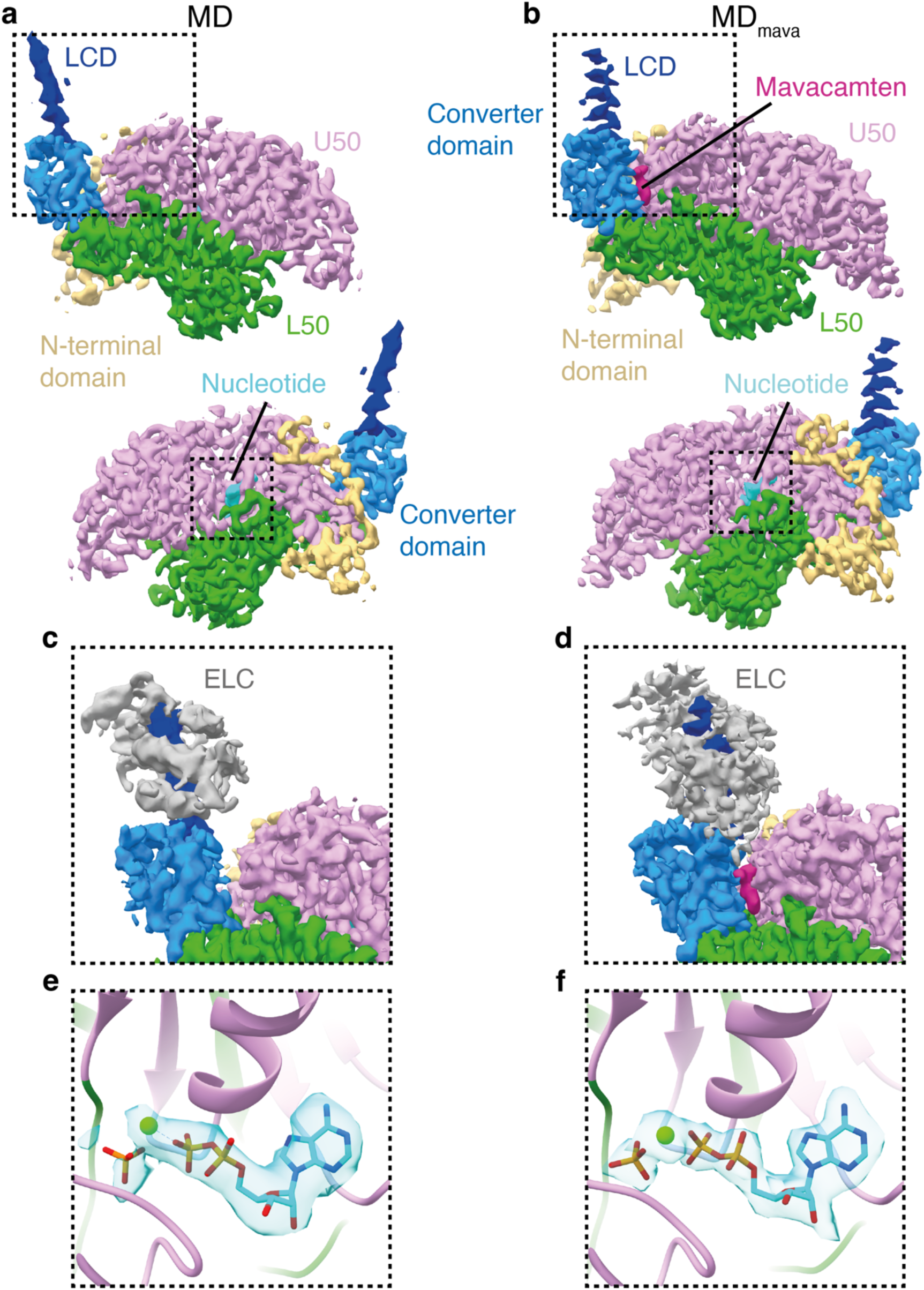
Primed motor domain cryoEM maps. (a) Segmented cryoEM map MD, split by subdomain (contour 0.5): N-terminal domain beige, L50 green, U50 pink, converter domain light blue and LCD dark blue. (b) Segmented cryoEM map of MD_mava_, split by subdomain (contour 0.6) coloured as (a) with mavacamten in burgundy. (c-d) Magnified view of segmented cryoEM map lever displaying ELC density grey (c) MD (contour 0.3) (d) MD_mava_ (contour 0.36) (e-f) ADP.Pi fit to segmented density colured by hetroatom (c) MD (contour 0.5) (d) MD_mava_ (contour 0.6).

**Extended Data Figure 5.**
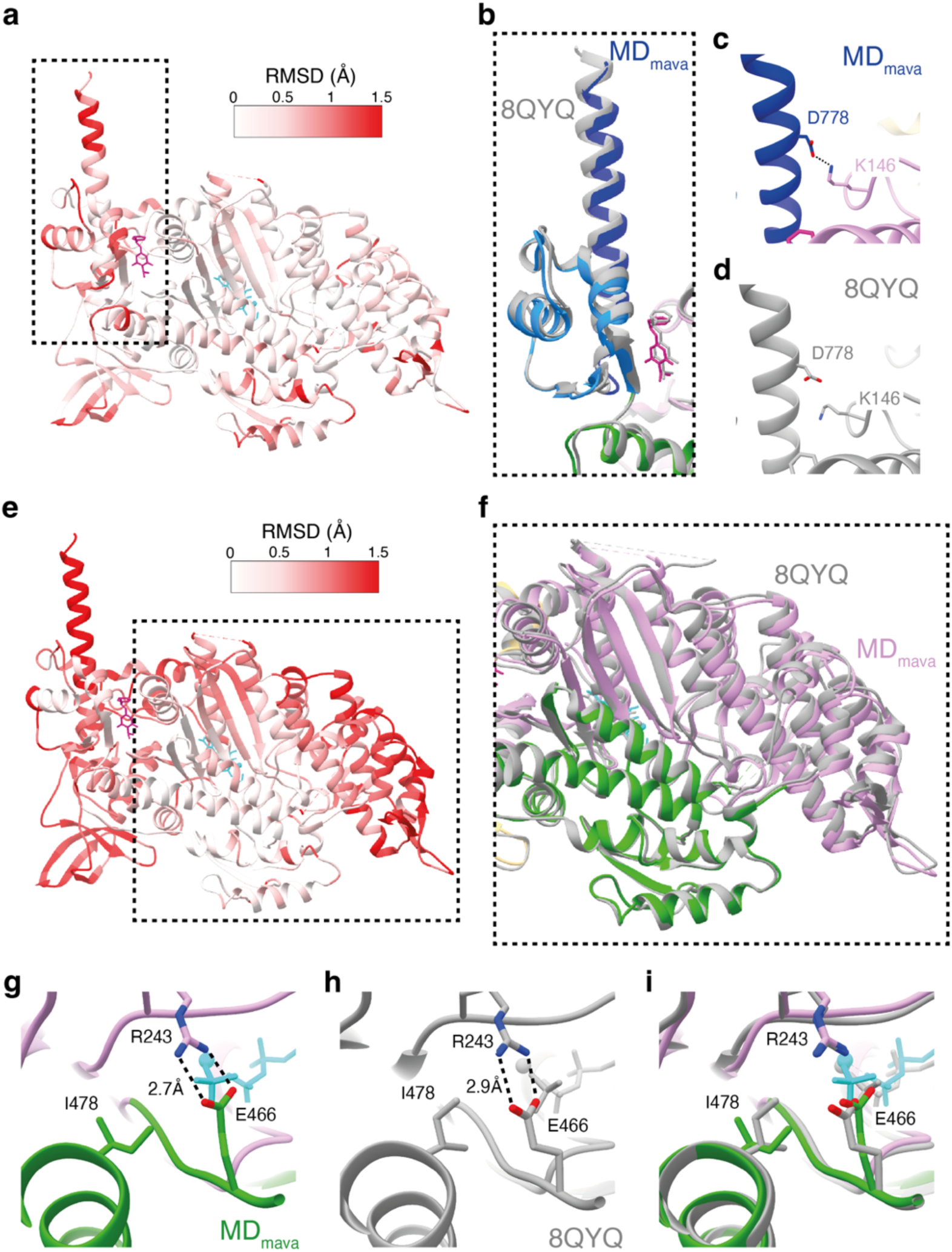
Comparison of MD_mava_ structure to bovine S1 crystal structure. (a) RMSD comparison between MD_mava_ and crystal structure of bovine S1 in complex with mavacamten (burgundy) (PDB ID: 8QYQ) coloured on MD_mava_ pdb (global structure alignment). (b) Overlay of lever position between MD_mava_, coloured by subdomain (L50 green, U50 pink, converter domain light blue, LCD dark blue and Mavacamten in burgundy) and 8QYQ grey (global structure alignment). (c-d) Magnified view of D778 and K146 interaction in (c) MD_mava_, coloured as in (b) and (d) 8QYQ grey. (e) RMSD comparison as in (a) but aligned using L50. (f) magnified view of (e) displayed as overlay of MD_mava_ coloured as in (b) and 8QYQ grey showing differing U50 conformation. (g-i) Magnified view of back door residues R243, E466 as well as I478 highlighting differing conformation between (g) MD_mava_ coloured as in (b) with the nucleotide in turquoise, (h) 8QYQ grey and (i) overlay.

**Extended Data Figure 6.**
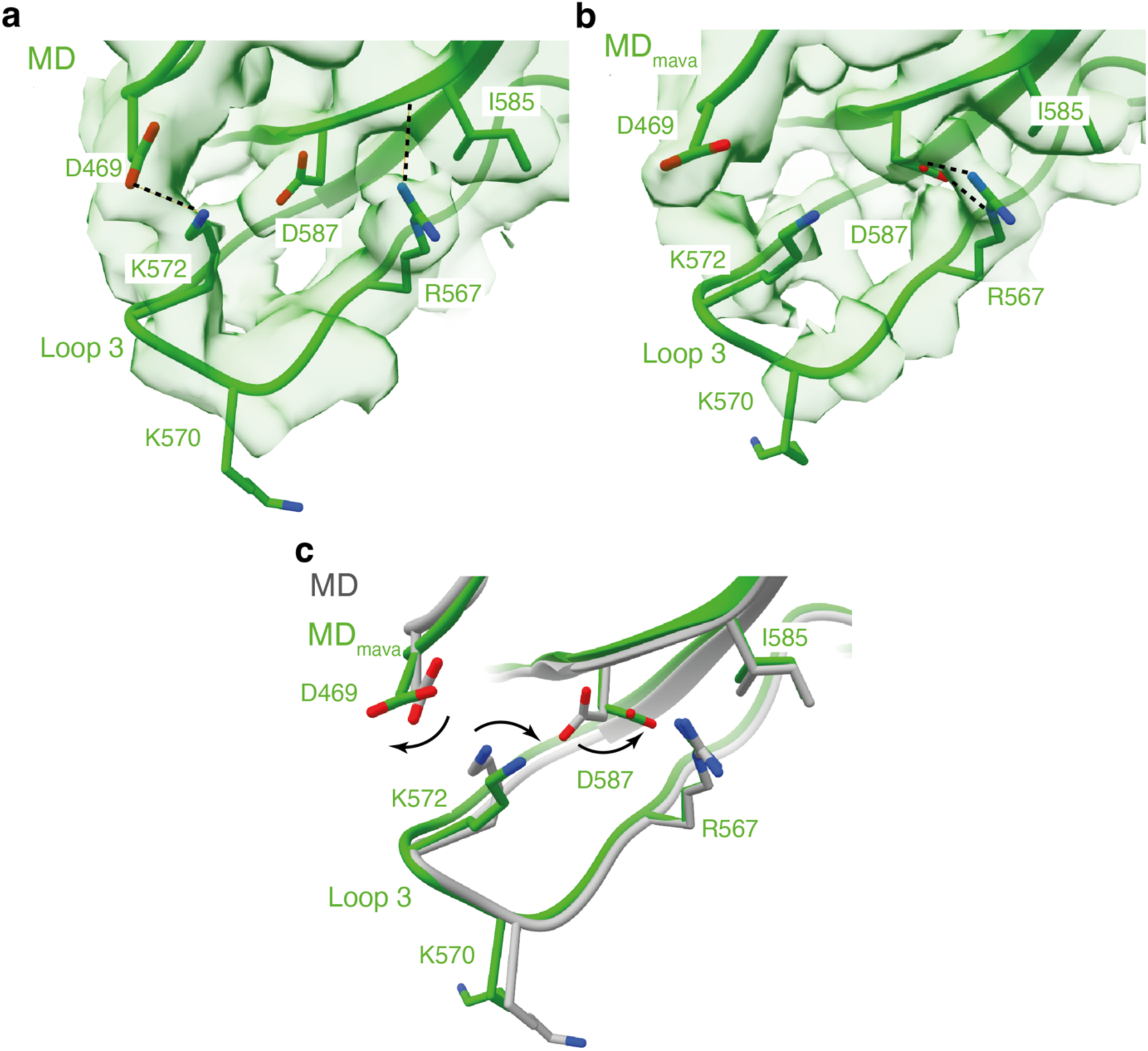
Allosteric effect of mavacamten on loop 3 hydrogen bonding. (a-c) Magnified view of loop 3 hydrogen bonding network. (a) MD model green in segmented cryoEM map (contour 0.25), highlighting hydrogen bonding between D469-K572 and R567-I585 alongside D587 position. (b) MD_mava_ model green in segmented cryoEM map (contour 0.42), highlighting hydrogen bonding between R567-D587 and D469, K572 position. (c) Overlay of MD gray and MD_mava_ green models highlighting change in D469, K572 and D587 resulting in loss of D469-K572 interaction explaining subsequent increase in crosslinking reactivity for K572 in the presence of mavacamten.

**Extended Data Figure 7.**
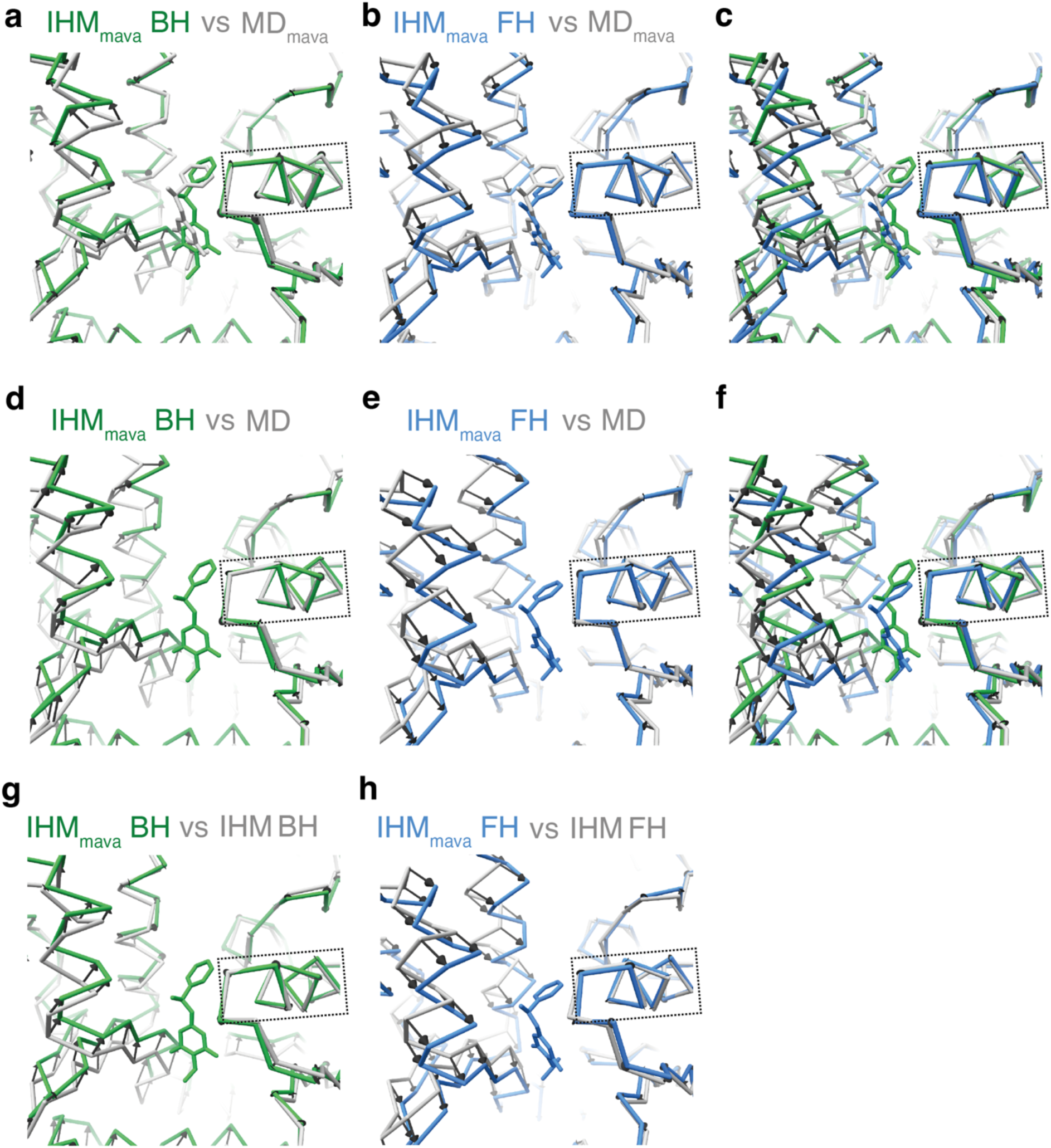
Change within the mavacamten binding site between the blocked and free head of the IHM. (a-c) Comparison of the MD_mava_ and IHM_mava_ mavacamten binding sites shown as backbone trace with vector arrows between alpha carbons. Models were aligned on the HE helix (residues 154-168) highlighted by the dashed box. (a) MD_mava_ grey and IHM_mava_ BH green. (b) MD_mava_ grey and IHM_mava_ FH blue. (c) Overlay of panels (a-b) to highlight that the conformational change is in opposite directions for the two heads of the IHM. (d-f) Comparison of the MD and IHM_mava_ mavacamten binding sites dispayed as in (a). (d) MD grey and IHM_mava_ BH green. (e) MD grey and IHM_mava_ FH blue. (f) Overlay of (d-e). (g-h) Comparison of the IHM and IHM_mava_ mavacamten binding sites displayed as in (a). (g) IHM BH grey and IHM_mava_ BH green. (h) IHM FH grey and IHM_mava_ FH blue.

**Extended Data Figure 8.**
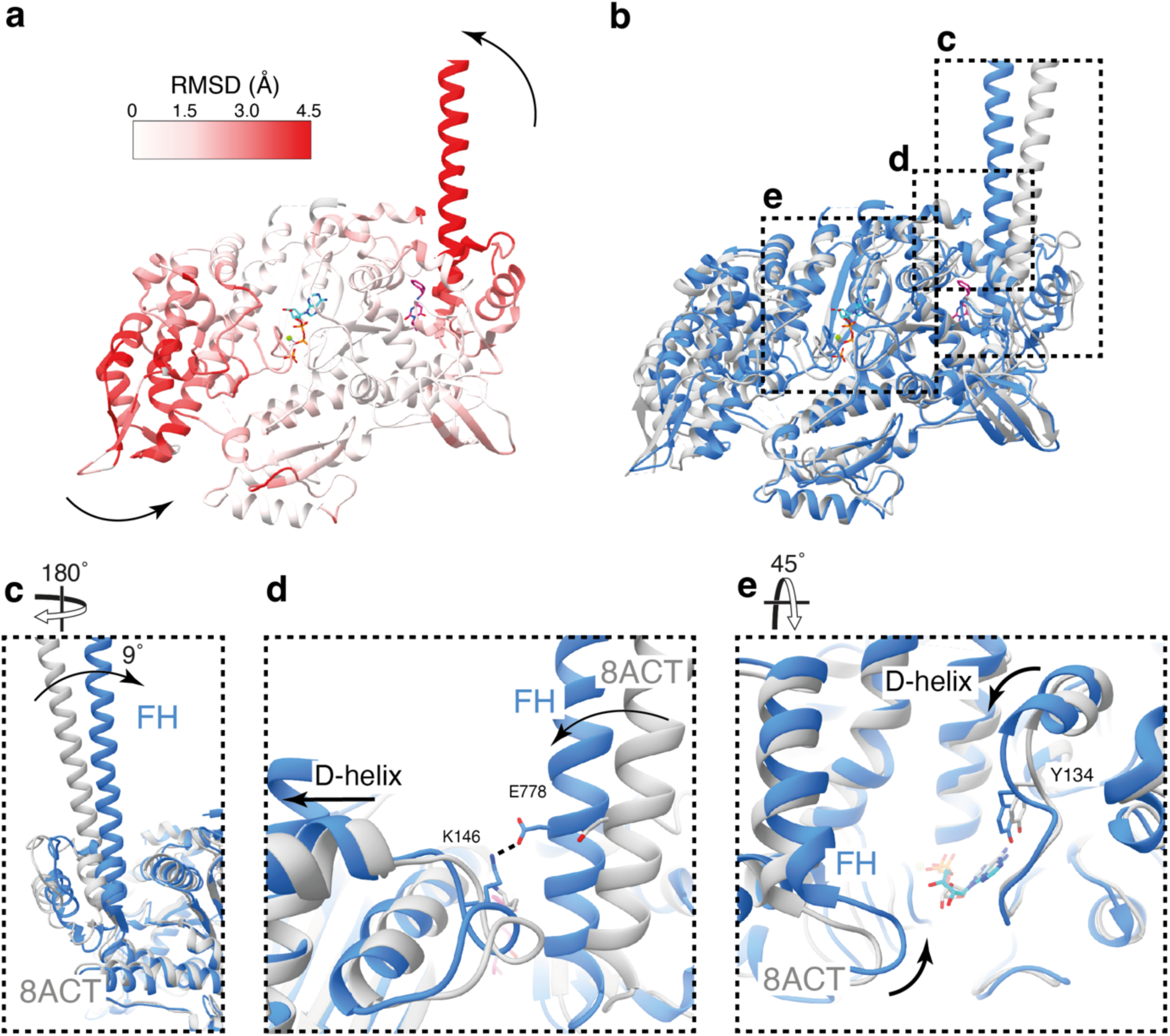
FH IHM_mava_ comparison to the IHM. (a) RMSD comparison between IHM_mava_ FH and folded-back state FH (PDB ID: 8ACT) aligned on the L50, coloured on IHM_mava_ FH model, highlighting domain movements. (b) Overlay of IHM_mava_ FH blue and folded-back state FH grey. (c) Side view of the IHM_mava_ FH lever overlaid on the folded-back state FH highlighting the 9° shift of the lever, coloured as in (b). (d) Structural comparison of lever and D-helix conformation showing E778-K146 coupling hydrogen bond in IHM_mava_ FH but not in the folded-back state FH, coloured as in (b). (e) Structural comparison of active site highlighting loop closure around active site in IHM_mava_, coloured as in (b).

**Extended Data Figure 9.**
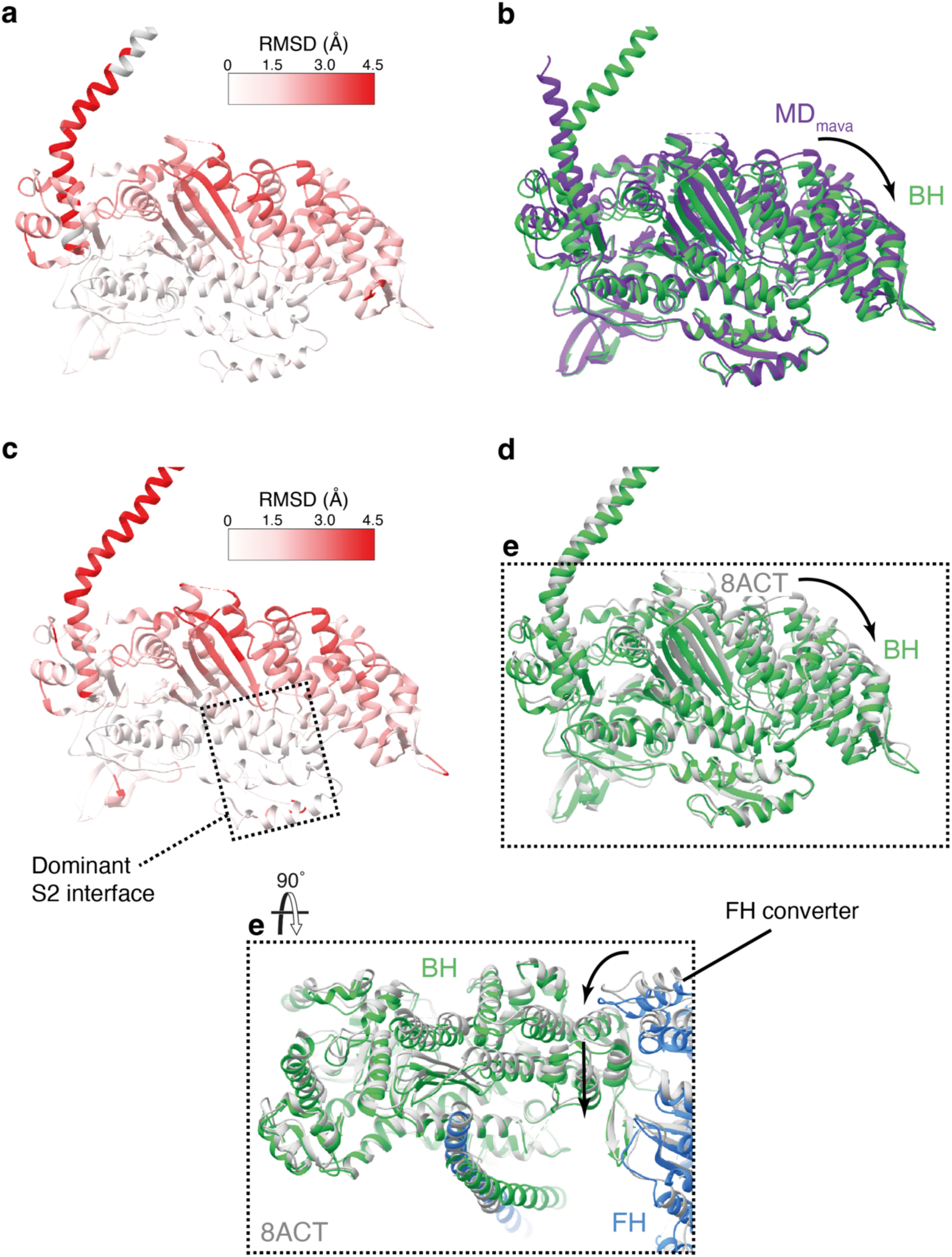
BH IHM_mava_ comparison to MD_mava_ and the IHM. (a) RMSD comparison between IHM_mava_ BH and MD_mava_ aligned on the L50, coloured on IHM_mava_ BH model (b) Overlay of IHM_mava_ BH green and MD_mava_ purple, highlighting U50 movement. (c) RMSD comparison between IHM_mava_ BH and folded-back state BH (PDB ID: 8ACT) aligned on the L50, coloured on IHM_mava_ FH model, highlighting the region the IHM_mava_ S2 has the most interactions with the BH in our model. (d) Overlay of IHM_mava_ BH (green) and folded-back state BH (grey), highlighting U50 movement. (e) Overlay of IHM_mava_ (green/blue) and folded-back state (grey) aligned on the BH, highlighting how the change in FH lever angle changes BH U50 conformation.

**Extended Data Figure 10.**
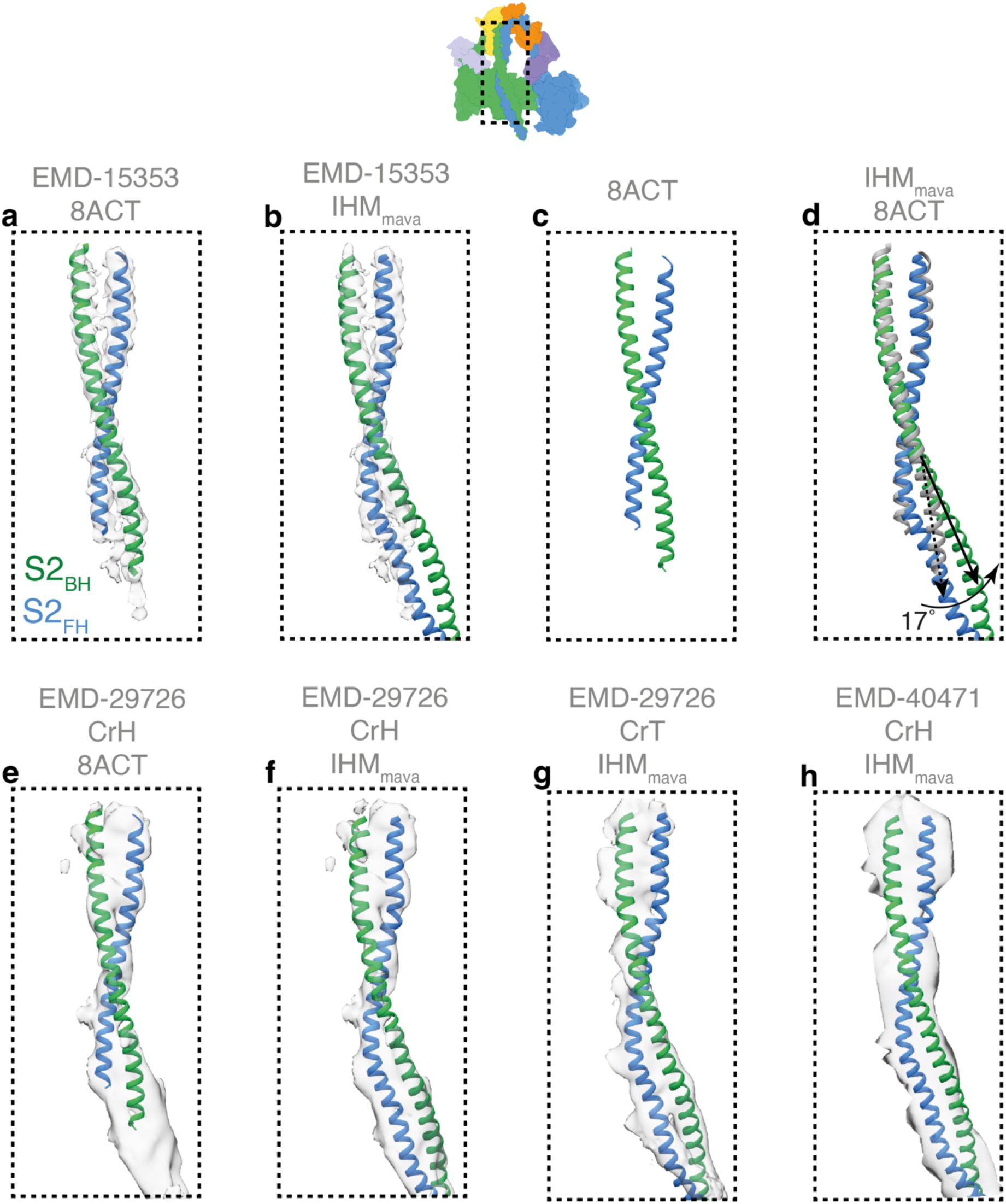
IHM_mava_ S2 comparison. (a-d) Comparison of S2 conformation between IHM_mava_ and IHM. (a) IHM S2 PDB ID: 8ACT (green/blue) rigidly fitted into corresponding segmented map EMD-15353 (contour 0.2). (b) IHM_mava_ (green/blue) rigidly fitted into segmented map EMD-15353 (contour 0.2). (c) IHM S2 model (green/blue). (d) IHM_mava_ S2 model (green/blue) compared to IHM S2 model (grey) highlighting BH heavy chain angle change of 17° (measured from BH 878-905). (e) Comparison of S2 conformation between IHM and mavacamten stabilised thick filament EMD-29726 horizontal crown (CrH)(contour 0.15) (f-h) Comparison of S2 conformation between IHM_mava_ and mavacamten stabilised thick filament EMD-29726 (f) horizontal crown (CrH)(contour 0.15) and (g) tilted crown (CrT)(contour 0.15).(h) Comparison of S2 conformation between IHM_mava_ mavacamten free thick filament CrH EMD-40471(contour 5.0).

**Supplementary Movie 1. Mavacamten reduces the number of moving filaments in cHMM and not cS1.** Movie demonstrates actin gliding movment displayed as summed plots of actin filament movement over 100 seconds of motility in Fig. 1b,c. Actin gliding is shown for cHMM and cS1 +/− 5 µM mavacamten, each panel correspond to 100 sec of movement captured at 5 frames/sec. Playback is at 20 fps (4x speed) to illustrate the slow movement at 5 µM mavacamten. Without drug the actin filaments glide smoothly over both cHMM and cS1 surfaces. However, at saturating [mavacamten] (5 µM) the fraction of moving filaments for cHMM decreases and movement is often interrupted by long pauses. Comparatively, filaments continue to move smoothly over the cS1 surface with only a reduction in speed.

**Supplementary Movie 2. Structural changes induced by mavacamten binding open motors.** Overview of key structural changes in the motor domain induced by mavacamten binding. Model state and morph direction is shown in the top left of the movie. (0:00) Overview of MD segmented cryoEM map (contour 0.5) split by subdomain: N-terminal domain beige, L50 green, U50 pink, converter domain light blue and LCD dark blue. (0:14) 360° rotation of MD PDB in segmented map. (0:30) Fade to un split sharpened MD cryoEM map (contour 0.56) and representation of mavacamten binding. (0:35) Morph of map and pdb from MD to MD_mava_. (0:44) Conformational changes induced by mavacamten binding, MD pdb grey and MD_mava_ pdb coloured. (0:58) 360° rotation of MD_mava_ pdb in segmented cryoEM map (contour 0.6) split by subdomain coloured as in (0:00) as well as Mavacamten in burgundy. (1:17) Magnified view of mavacamten binding site. (1:35) magnified view of LCD D-helix interaction. (1:39) Fade to un split sharpened MD_mava_ cryoEM map (contour 0.56) followed by morph of map and model from MD_mava_ to MD and back. (1:51) Highlighting conformational change at the D-helix between MD grey and MD_mava_ coloured. (2:17) Magnified view of MD_mava_ back door pdb in segmented cryoEM map shown as in (0:58). (2:20) Fade to un split sharpened MD_mava_ cryoEM map (contour 0.56) followed by morph of map and model from MD_mava_ to MD and back. (2:34) Highlighting conformational change at the back door between MD grey and MD_mava_ coloured.

**Supplementary Movie 3. IHM_mava_ interaction interfaces.** (0:00) Overview and 360° rotation of IHM_mava_ segmented cryoEM map coloured by chain (contour: 0.08): blocked head green, free head blue, blocked head ELC light purple, free head ELC purple, blocked head RLC yellow, free head RLC orange, mavacamten burgundy and the nucleotide in light blue. (0:31) 360° rotation of IHM_mava_ PDB in segmented cryoEM map coloured as in (0:00). (0:50) Magnified view of BH mavacamten binding site. (1:07) Magnified view of FH mavacamten binding site. (1:30) Magnified view of motor-motor interface (change in map contour to 0.01). (1:46) Magnified view of HCM loop_BH_ transducer_FH_ interface. (2:03) Magnified view of BH ELC_FH_ interface. (2:22) Overview of S2 BH interface highlighting the three main contact regions on the BH: OH-helix, W-helix and HLH. (2:36) Magnified view of S2 OH-helix_BH_ interface. (2:49) Magnified view of S2 W-helix_BH_ interface. (3:01) Magnified view of S2 HLH_BH_ interface.

