## Supplementary Information for "Mavacamten inhibits myosin activity by stabilising the myosin interacting-heads motif and stalling motor force generation"

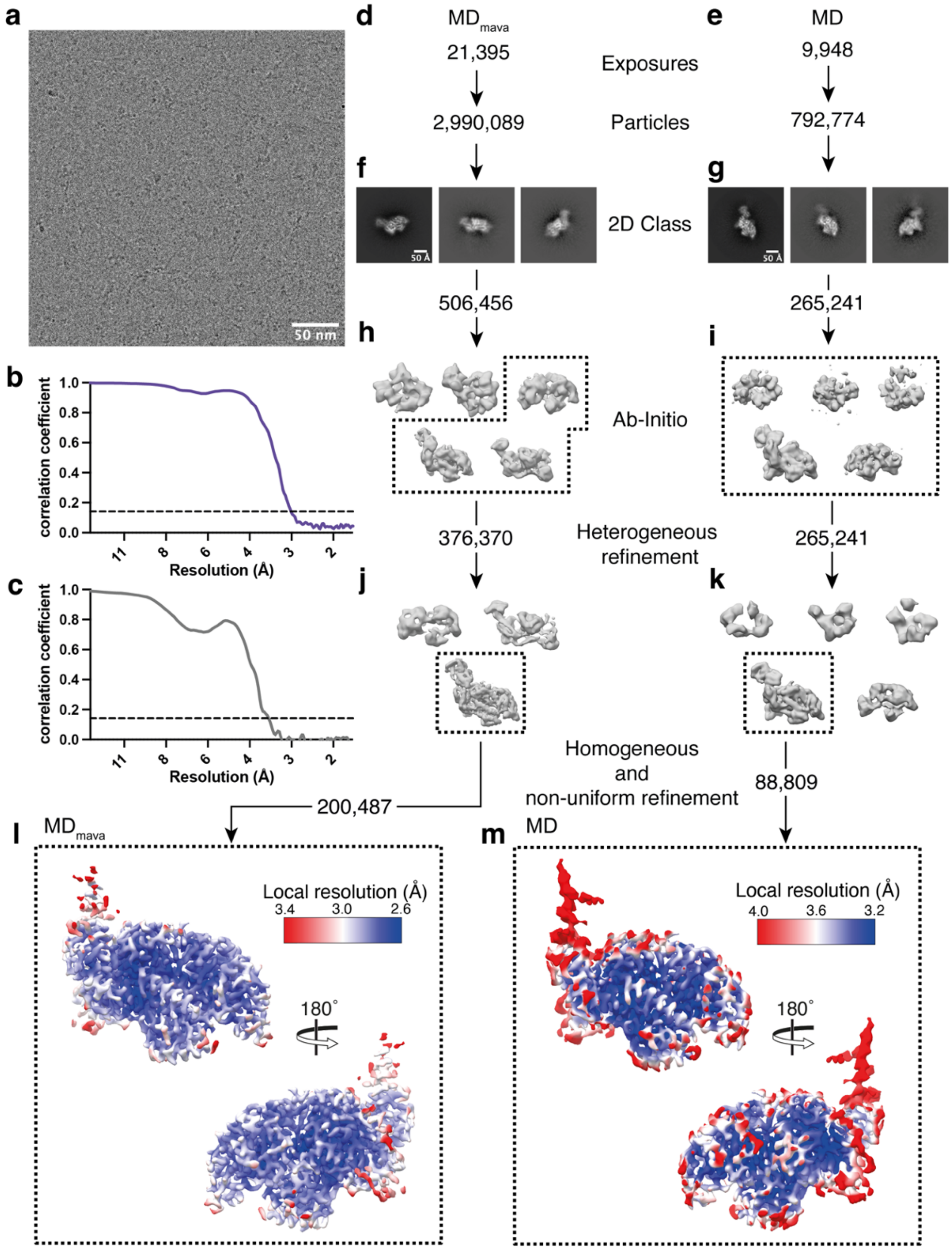


**Supplementary Figure 1: Open head motor domain processing pipeline.** (a) Representative cryoEM micrograph from $MD_{mava}$ dataset. (b-c) FSC curves, $MD_{mava}$ 2.9Å and MD 3.4Å respectively, 0.143 threshold represented by dashed line. (d-m) Flow diagram for $MD_{mava}$ and MD data processing. (d,e) Number for micrographs collected and particles extracted for $MD_{mava}$ and MD respectively (f,g) Representative 2D classes. (h,i) Ab-initio classes, selected classes carried forward for refinement are boxed. (j,k) heterogeneous refinement classes. (l,m) $MD_{mava}$ and MD final reconstruction respectively, coloured by local resolution estimation.

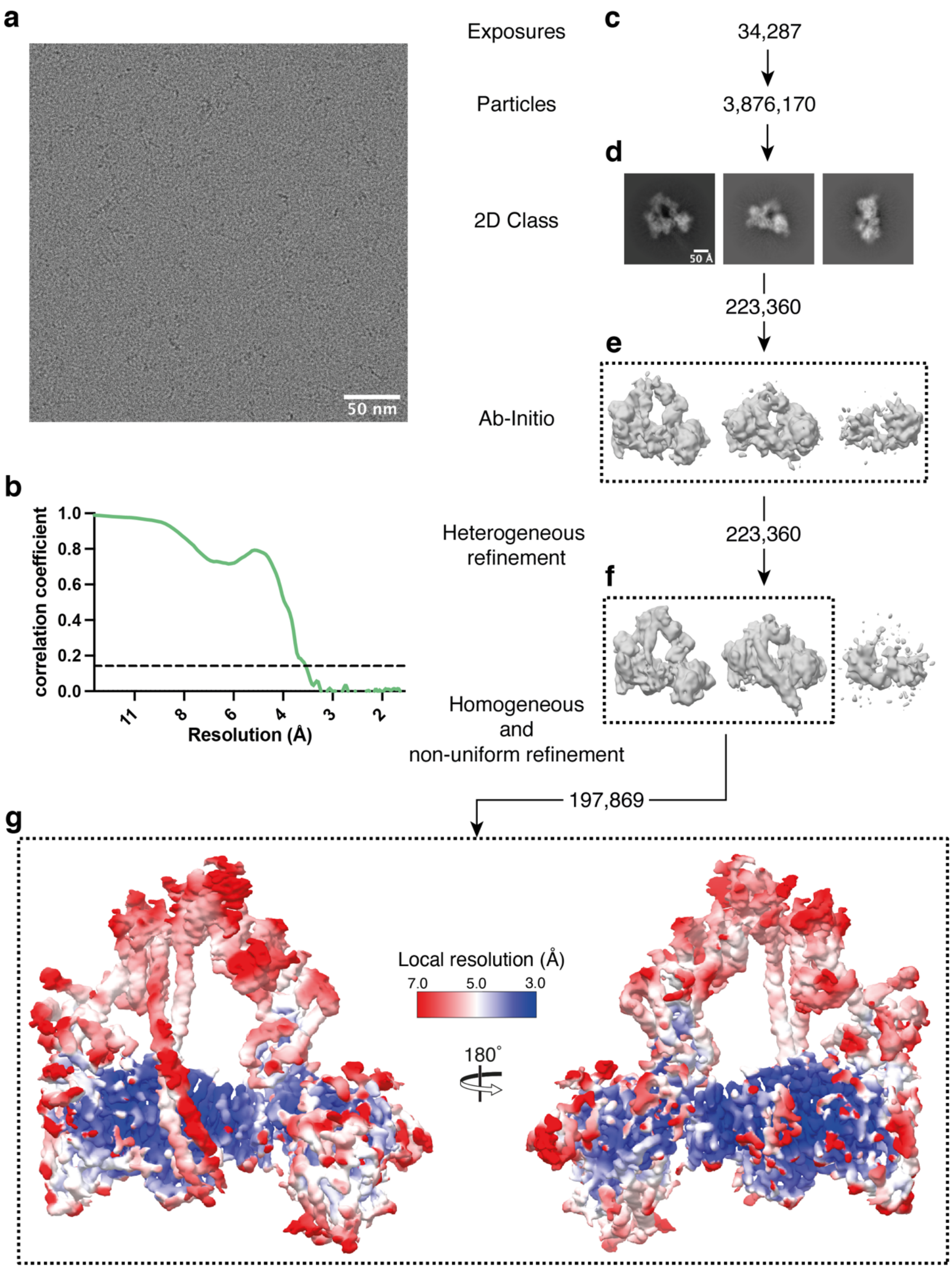


**Supplementary Figure 2: IHM$_{mava}$ processing pipeline.** (a) Representative cryoEM micrograph from IHM$_{mava}$ dataset. (b) IHM$_{mava}$ FSC curve 3.7Å, 0.143 threshold represented by dashed line. (c-g) Flow diagram for IHM$_{mava}$ image processing. (c) Number of micrographs collected and particles extracted for IHM$_{mava}$. (d) Representative 2D classes. (e) Ab-initio classes, selected classes carried forward for refinement are boxed. (f) Heterogeneous refinement classes. (e) IHM$_{mava}$ final reconstruction, coloured by local resolution estimation.

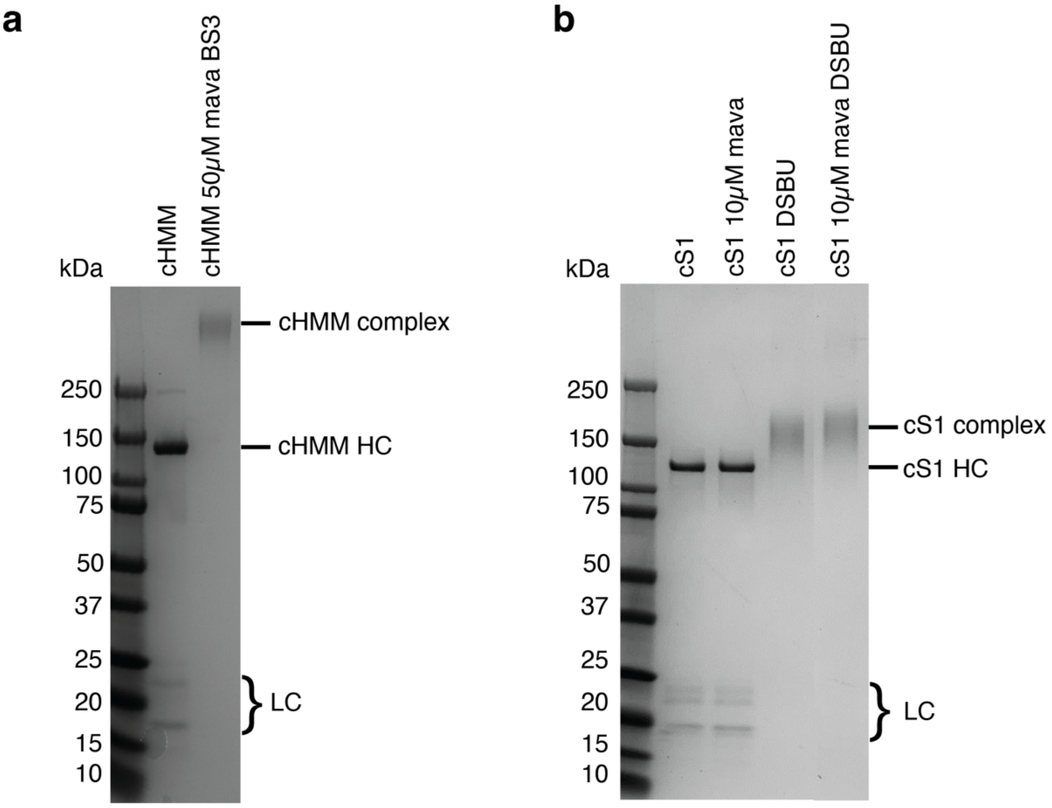


**Supplementary Figure 3: SDS-PAGE gel of crosslinked cHMM and cS1 complexes.** All gels shown 4–20% Mini-PROTEAN® TGX™ Precast Protein Gels, 15-well (BioRad) and precision plus dual colour standards (Biorad) ladder (a) cHMM un-crosslinked and BS3 crosslinked samples highlighting the crosslinked cHMM complex, cHMM heavy chain (HC) and associated light chains (LC). (b) cS1 un-crosslinked and DSBU crosslinked samples highlighting the cS1 complex, HC and LCs.
